# Allele mining, evolutionary genetic analysis of *TaHKT1;5* gene and evaluation of salinity stress in selected lines of wheat

**DOI:** 10.1101/2022.04.01.486792

**Authors:** Karthikeyan Thiyagarajan, Prem Narain Mathur, Prashant Vikram, Deepmala Sehgal, Ravi Valluru, Velu Govindan, Hifzur Rahman, Dhruba Bahadur Thapa, Sumitra Pantha, Patrizia Galeffi, Arianna Latini, Cristina Cantale, Enrico Porceddu, Arun Kumar Joshi

**Affiliations:** Bioversity International, Rome, Italy; International Maize and Wheat Improvement Center (CIMMYT), El Batan, Mexico; Borlaug Institute for South Asia (BISA)-CIMMYT, New Delhi, India; University of Linclon, Lincoln, United Kingdom; Nepal Agricultural Research Council (NARC), Lalitpur, Nepal; Italian National Agency for New Technologies, Energy and Sustainable Economic Development-ENEA, Rome, Italy; Accademia Nazionale dei Lincei, Rome, Italy; International Center for Biosaline Agriculture, Academic City, Dubai, United Arab Emirates

**Keywords:** *High Affinity Potassium Transporter* (HKT), *HKT* gene, *Triticum*, Diversity, Evolution, Phylogenetics, Phenotype, Salinity stress, Ditelosomic lines, SNPs, Transposable Elements, Allelic Variant

## Abstract

This study reports the novel allelic diversity in *HKT 1;5* gene (*High Affinity Potassium Transporter*) in bread wheat and its phylogenetic relationship among the paralogs/orthologs of in *Triticum aestivum* and its wild relatives. *HKT 1;5* gene is a known and pivotal gene associated with the salinity tolerance in plants upon the discrimination of K^+^ over Na^+^ in leaves without change in Na^+^ concentration in root. This gene was sequenced in a diverse collection of bread wheat, durum wheat, wild relatives, and ditelosomic lines. Sequence analysis in bread wheat led to the identification of four alleles, which could be distinguished by number of SNPs. Sequence comparison between monocot bread wheat and dicot *Arabidopsis thaliana* revealed that the *HKT1;5* gene is conserved at level of exonic regions; however, the presence of transposable elements especially in intronic regions is further intriguing towards evolutionary relatedness. Two paralogous or major alleles observed in *Triticum monoccum* and *Aegilops tauschii* were further categorized as sub-alleles based on their SNPs comparison. This gene was absent in *T. urartu* in accordance with existing evidence, while it was found in *A. speltoides* (an allelic variant) with a few base pairs insertion in the exon1 region causing a frameshift mutation with an altered amino acids and genomic database mining unveiled additional alleles in this species. Ditelosomic lines with 4DL and 4DS chromosomes revealed a higher similarity with bread and durum wheat respectively. Phylogenetic studies of *HKT1;5* orthologs from different Poaceae species revealed the occurrence of five different ortholog groups with taxonomic consistency. Phenotyping based salinity stress experiment distinguished the unknown lines for salinity tolerance and sensitiveness in comparison with known reference lines and possible allelic comparison was made. The salinity stress analysis further revealed that some known drought/heat tolerance lines showed slightly better salinity tolerance with mean values and variability of traits than known saline tolerant wheat line at controlled ambient.

## INTRODUCTION

Salinity or sodicity marks 6% of the world’s land covering around 400 million hectares of cultivated and uncultivated land (<u>FAO Land and Plant Nutrition Management Service, 2008</u>). Soil salinity imposes ion toxicity, osmotic stress, nutrient (N, Ca, K, P, Fe, Zn) deficiency and oxidative stress on plants. It is one of the main abiotic stresses resulting in the reduction of growth and yield of different crops including wheat (Rengasamy, 2010). In several wheat-producing countries of the Indian subcontinent (e.g. India and Pakistan) and the Middle Eastern region (Iran, Egypt, and Libya), salinity affects up to 10% of the total wheat belt, but in some regions, such as Western Australia, salinization affects more than half of wheat farms (Colmer et al. 2006, Honghong et al. 2014). Salt concentrations beyond 40 mM or electrical conductivity greater than 4 dS/m in soil has been reported to determine significant reduction in number of spikelets per spike, delayed spike emergence and reduced fertility, resulting in poor grain yields (Shrivastava and Kumar 2015, Kalhoro et al., 2016). Adverse of effect of salinity during developmental stages of wheat and strategies to overcome problems associated with salinity stress has been recently reviewed (Sabagh et al., 2021). The role of salinity tolerance concerning *HKT1;5* gene and its translocation events were shown based on the available research evidence (James et al., 2011, Byrt et al., 2007) (Fig. 1A, 1B). Salinity tolerance and causes of sensitivity of plants is depicted with an illustration (Fig. 1C).

**Figure 1.**
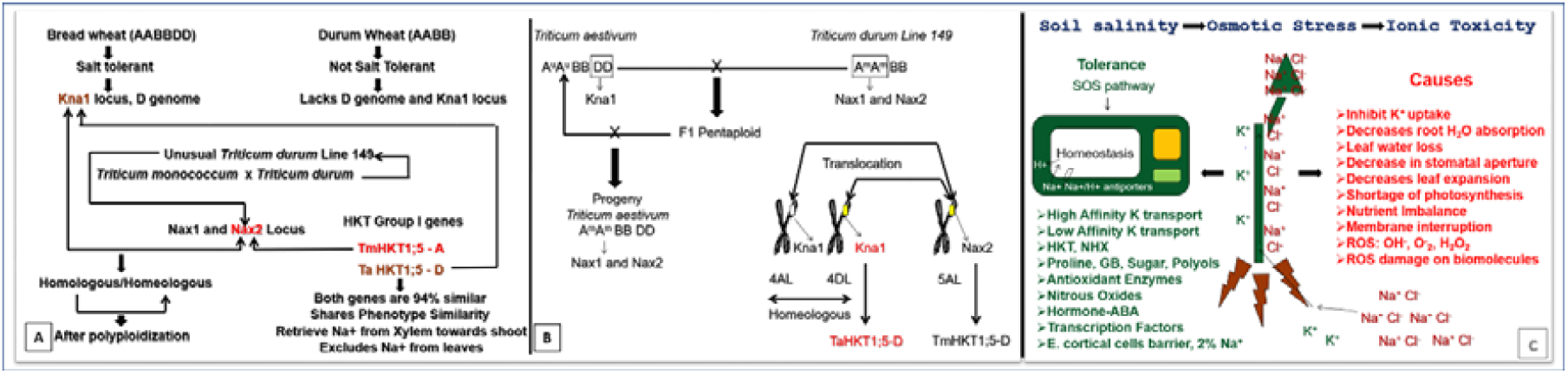
A). Genes behind the salinity tolerance in wheat. B). Reciprocal translocation of *Kna1* and *Nax2* locus. Schematic representation. Based on sources: James et al., 2011, Byrt et al., 2007. C) Schematic representation of salinity tolerance and sensitiveness in plants.

Improving salinity tolerance is an important breeding target in wheat breeding programs. A major genetic mechanism contributing towards salinity tolerance of most crops including the cereals rice (*Oryza sativa*) and wheat (*Triticum aestivum* and *Triticum monococcum*) is Na^+^ exclusion from leaves. High-affinity potassium (K^+^) transporter (*HKT*) protein family have been reported to play a major role in this trait (Ren *et al*. 2005; James *et al*. 2006; Munns *et al*. 2012; Munns and Gilliham, 2015; Campbell *et al*. 2017; Xu *et al*. 2018). Several *HKT* genes, including *HKT1;1*-like, *HKT1;3*-like, *HKT1;4*-like, and *HKT1;5*-like, have been identified and mapped to wheat homoeologous chromosome groups 2, 6, 2, and 4, respectively (Huang et al. 2008). *HKT 1;5* gene associated *Kna1* locus was reported for salinity tolerance in plants due to the discrimination of K^+^ over Na^+^ in leaves with net Na^+^ loading in root xylem than net uptake (Gorham et al., 1990). The wheat sodium transporters *TmHKT1;5-A* and *TaHKT1;5-D* (Genbank: DQ646342) are encoded by genes underlying major shoot Na^+^ exclusion loci *Nax2* (chromosomal location: 5AL, ancestrally 4AL) and *Kna1* (chromosomal location: 4DL) from *Triticum monococcum* (Tm) and *Triticum aestivum* (Ta), respectively (Byrt et al., 2007). Role of *TmHKT1;5-A* in improving grain yield on saline soils by 25% has been reported (Munns et al 2012). The presence of *HKT* gene in several plant species has been reported (Waters e al., 2013). Comparative genomics suggested that there is no synteny of the *HKT1;5* gene between *O. sativa* and bread wheat unlike other *HKT* genes (Huang *et al*., 2008), While only a single copy of the *HKT* gene is (Uozumi *et al*., 2000) exist in model dicot species *A. thaliana*, which further supported by the *HKT* gene search on *A. thaliana* genome (Mayer *et al*., 1999). The divergence of this gene must have occurred early during evolution of monocots and dicots (Waters et al., 2013). Allele mining pertain to natural variations is useful strategy for the identification of better salinity tolerant variety from natural populations (Baxter et al., 2010). Efforts have taken to genetically characterize the *HKT1;5* gene to better understand the molecular evolutionary basis of diversity of salt tolerance among bread wheat and its wild relatives. Therefore, understanding the allelic diversity, natural variations, and divergence of *HKT1;5* gene would be an asset and provides the genetic basis of the evolution of this gene. In this study, using a diverse wheat material and multifaceted genetic approaches, we demonstrate that *HKT1;5* gene has multiple allelic variants present within the wheat germplasm pool and its wild progenitor species.

## MATERIALS AND METHOD

### Plant materials and DNA extraction

The plant materials included 16 bread wheat and one durum wheat cultivars, and two ditelosomic lines (DTGH09_1559, possessing 4DL and lacking 4DS chromosome, and DTGH09_1556, possessing 4DS and lacking 4DL chromosome). Furthermore, a total of 18 wild accessions were selected to represent the diploid progenitors of the three genomes of bread wheat, namely *Triticum monococcum* (Am genome), *Triticum uratu* (Au genome), *Aegilops speltoides* (B genome) and *Aegilops tauschii* (Dt genome). After attaining four leaf stage, DNA was extracted from collected leaves of each cultivar using the CTAB method (Murray and Thompson, 1980) (Table 1). After attaining four leaf stage, DNA was extracted from collected leaves of each cultivar using the CTAB method (Murray and Thompson, 1980).

**Table 1.** Saline tolerant/sensitive, unknown bread wheat lines, a durum durum wheat, ditelosomic lines and wheat wild progenitors/relatives

| No. | Variety | Country | Source | Species |
| --- | --- | --- | --- | --- |
| 1 | Kharchia | India | CIMMYT | *T. aestivum* |
| 2 | Kharchia 65 | India | CIMMYT | *T. aestivum* |
| 3 | Shorawaki | Pakistan | CIMMYT | *T. aestivum* |
| 4 | Chinese Spring | China | CIMMYT | *T. aestivum* |
| 5 | YMI#6 | China | CIMMYT | *T. aestivum* |
| 6 | Pasban 90 | Pakistan | CIMMYT | *T. aestivum* |
| 7 | KRL 1-4 | India | CIMMYT | *T. aestivum* |
| 8 | Lu 26-S | Pakistan | CIMMYT | *T. aestivum* |
| 9 | Sakha 8 | Egypt | CIMMYT | *T. aestivum* |
| 10 | Oasis F86 | Mexico | CIMMYT | *T. aestivum* |
| 11 | C306 | India | CIMMYT | *T. aestivum* |
| 12 | Pastor | Mexico | CIMMYT | *T. aestivum* |
| 13 | Munal#1 | Nepal | NARC | *T. aestivum* |
| 14 | DharwarDry | India | NARC | *T.aestivum* |
| 15 | HDR-77 | India | NARC | *T.aestivum* |
| 16 | WK1204 | Nepal | NARC | *T.aestivum* |
| 17 | DWK26 | Nepal | NARC | *T.durum* |
| 18 | DTGH09_1559 | Mexico | CIMMYT | *T.aestivum_Dt* |
| 19 | DTGH09_1556 | Mexico | CIMMYT | *T.aestivum_Dt* |
| No. | ID | Entry | Source | Species |
| 20 | PI272558 | 701 | CIMMYT | *T. monococcum* |
| 21 | PI428150 | 733 | CIMMYT | *T. monococcum* |
| 22 | PI326317 | 716 | CIMMYT | *T. monococcum* |
| 23 | PI264935 | 700 | CIMMYT | *T. monococcum* |
| 24 | PI272560 | 702 | CIMMYT | *T. monococcum* |
| 25 | PI428333 | 171 | CIMMYT | *T. urartu* |
| 26 | PI428335 | 172 | CIMMYT | *T. urartu* |
| 27 | PI428258 | 188 | CIMMYT | *T. urartu* |
| 28 | PI487265 | 173 | CIMMYT | *T. urartu* |
| 29 | PI428257 | 187 | CIMMYT | *T. urartu* |
| 30 | CWI94200SH(125) | 23 | CIMMYT | *A. speltoides* |
| 31 | CWI94201SH(126) | 24 | CIMMYT | *A. speltoides* |
| 32 | CWI94203SH(128) | 25 | CIMMYT | *A. speltoides* |
| 33 | CWI94205SH(130) | 26 | CIMMYT | *A. speltoides* |
| 34 | CWI94207SH(132) | 27 | CIMMYT | *A. speltoides* |
| 35 | CWI94208SH(133) | 28 | CIMMYT | *A. speltoides* |
| 36 | CWI94958SH(1344) | 443 | CIMMYT | *A. tauschii* |
| 37 | CWI94956SH(1342) | 441 | CIMMYT | *A. tauschii* |

### NCBI GenBank Submission and retrieval of sequences from public databases for Phylogeny

So far, we have generated and used 29 plausible nucleotide sequences; among them, 8 were about 3 kb long and were complete coding sequences exclusively from wheat. The remaining 21 sequences were about 1 kb long and are from different progenitor species, ditelosomic lines and some varieties of wheat as well.

For comparative and phylogenetic analysis (besides the sequences generated in this study), additional sequences of *HKT1;5* gene were retrieved from Public Data Bases such as NCBI, URGI-IWGSC etc. Concerning the alleles of bread wheat lines, the genomic positions of the sequences and SNPs were compiled at ENSEMBL-Wheat database server and SNPs extraction was done using Linux based snp-sites program (Page et al., 2016). SNPs extracted fasta file was used as input to create Hapmap and VCF (Variant Call Format) files in Tassel (Bradbury et al., 2007). Subsequently, SIFT-Sorting Intolerant from Tolerant (Sim et al., 2012) analysis was carried out using SNPs derived VCF file of *T. aestivum HKT1;5* at ENSEMBL Wheat database’s VEP server and further comparisons were made using TraesCS4D02G361300 from ENSEMBL. *A. thaliana HKT1* gene was derived from Chromosome 4 (NCBI: CP002687, Mayer et al., 1999) and confirmed with Uozumi et al., 2000.

### Primer design, Polymerase Chain Reaction and Sequencing

A and D genome specific primers for the *T. aestivum HKT1;5-D* and *T. monococcum HKT1;5* genes were designed using the reference sequences available at NCBI (Genbank: DQ646342 DQ646339, Byrt *et al*., 2007). Primers allowed amplifying the gene from start to stop codon within a single Polymerase Chain Reaction (PCR) in genome specific manner. Alternatively, additional UTR anchoring, and internal primers were used to amplify different gene fragments that overlapped to obtain the full-length gene (Supplementary Table 1). The primers designed from *T. monococcum* were also amplified a gene in *A. tauschii* and *T. aestivum* 4DL or *A. tauschii* specific primers were also amplified a gene in *T. monococcum*. The PCR amplification was performed in 20 µl reaction volume consisting of 1 µl of 50 ng template DNA, 0.5 µl of dNTP (2.0 mM each), 0.1 µl of Phusion™ Hifidelity DNA Polymerase (2U/µl) (Thermo Fisher Scientific Co.), 0.4 µl of 1 pM forward primer, 0.4 µl of 1 pM reverse primer, 0.6 µl of DMSO (100%), 4 µl of HF5X buffer, 0.4 µl of MgCl_2_ (50 mM), and 12.6 µl of Sigma H_2_O. PCR was performed using Mastercycler^®^ pro (Eppendorf) using the following cycling program: Initial denaturation at 98 °C for 40 s, 35 cycles of 12 s denaturation at 98 °C, 45 s annealing at 64 °C, 1.4 min extension at 72 °C and final extension at 72 °C for 5 min. PCR conditions and cycles were adjusted based on primer pair, amplicon length, GC content and annealing temperature etc. Subsequently, amplified PCR fragments were resolved on 1.2% agarose gel pre-stained with GelRed™ (Biotium Inc.) at 90 V for 90 min. PCR fragments were visualized using a UV illumination-based documentation unit. The visualized bands were cut from the agarose gels using clean scalpel and purified using Sigma Aldrich GenElute agarose gel purification kit. Additionally, amplified fragments were purified using a Qiagen QIAquick PCR purification kit (Cat. No. 28106) following the manufacturer’s instructions. The purified PCR fragments were sequenced through Macrogen Co., South Korea.

### Identification of SNPs for allele mining

The quality of chromatograms was reviewed based on PHRED quality scores using FinchTV (Geospiza Inc., USA) chromatogram viewer software. Chromatogram ab1 extension files were assembled using reference sequence (Byrt et al., 2007) with CodonCode aligner program (https://www.codoncode.com/aligner/). Alternatively, clear chromatogram regions were copied as text and converted to FASTA format as an input for the CAP3 assembly program and assembled to display the entire gene (Huang and Madan, 1999). Subsequent sequence alignment of the gene from different cultivars was performed using Clustal Omega with default parameters (Sievers *et al*., 2011). The presence of SNPs was manually detected with multiple sequence alignment, based on the alignment SNPs derived different alleles were designated. Whole gene from selected lines of bread wheat was sequenced and used for SNPs/allele prediction and relevant analysis, while in durum wheat, wild progenitors and ditelosomics lines, region spanning around 1 Kb at 5’ UTR and/or exon1 was exclusively sequenced and focused to detect UTR or coding sequence variations and for related studies.

### Identification of Repeated Elements

A search for identification of transposon and its related repeated elements in whole *HKT* gene of *T. aestivum* and *A. thaliana* was done using Repbase Censor server (https://www.girinst.org/censor/index.php). A predicted complete non-autonomous mariner transposon element was subjected for BLASTN analysis with IWGSC-URGI server (https://wheat-urgi.versailles.inra.fr/Seq-Repository/BLAST).

### Phylogenetic analysis

The sequences were aligned using Clustal Omega (Sievers *et al*., 2011) or Clustal X and exported as multiple sequences in aligned FASTA format (Larkin et al., 2007). Phylogenetic analysis of the gene sequences and relative divergence using RelTime (Mello et al., 2018) were carried out with MEGA11 (Molecular Evolutionary Genetic Analysis software) and gaps/missing parameters were excluded (Tamura et al., 2021). The Maximum Likelihood method with Jukes Cantor (JC) model and 1000 bootstrap resampling was used to reconstruct the phylogenetic tree of the sequences generated in this study along with the reference and orthologous gene sequences retrieved from public databases. Global alignment of the coding sequences was done using LAGAN alignment toolkit (Brudno *et al*., 2003) to validate the exonic conservation of *T. aestivum* and *A. thaliana*. Genetic analysis was done using DNAsp for Gamma statistics for gene flow estimates of haplotypes (Nei 1973), DeltaST (Nei, 1982), Nst (Lynch, 1990), Fst (Hudson *et al*., 1992) (Rozas J, 2017).

### *In silico* analysis of protein and 3-D modeling

Corresponding putative proteins of *A. thaliana HKT1* gene and *T. aestivum HKT1;5-D* gene tertiary structure protein modeling was done using HHPhred (Söding *et al*., 2005) using protein templates from Protein Data Bank (http://www.rcsb.org/pdb/home/home.do). Visualization and annotation of generated protein models were done with Rasmol (Sayle and Milner-White, 1995) and compared with the primary structure of the protein. Secondary structure was predicted using HNN secondary structure prediction method (Guermeur, 1997). Documentation of protein physico-chemical parameters including instability index was calculated using Protparam tool at Expasy server http://web.expasy.org/tools/protparam/protparam-doc.html (Gasteiger *et al*., 2005).

### PETRIPLATE BASED PHENOTYPIC TRAITS MEASUREMENT AND ANALYSIS

#### Completely randomized design experiment

Seeds of the bread wheat cultivars C306, HDR-77, Oasis F86 and WK1204 were surface sterilized with 5% of sodium hypochlorite for 5 minutes and were subsequently exposed to five salinity treatments: Control (Water), 50 mM, 100 mM, 150 mM, 200 mM salt solution. For each cultivar, 15 seeds were used for each of the five treatments in each experiment including control. The 15 seeds were placed between the layers of cellulosic filter papers. During the experiment, Petri plates were covered with lid to reduce the excess evaporation and for maintaining homogenous ambience. The average temperature of the experimental condition was 25.13°C (range 23-27.8°C) during the day and 14.29°C (range 12-18°C) during the night, with 8 hours of photoperiod during day. Salinity or control treatment was performed with three replicates through a completely randomized design. To compensate the evaporation, an equal amount of water was given to all samples periodically.

#### Phenotypic measurements: estimation of germination traits

Phenotypic traits viz. Final Germination Percentage (FGP), Germination Energy Percentage (GEP), Radical Length (RL), Number of Radicles (NoR), Coleoptile Length (CL), Relative Water Content (RWC), Fresh Weight (FW) and Dry Weight (DW) were measured. Germination was counted at every one-day interval until the 7^th^ day when the germination was ceased. Each day was assigned as T for a day, i.e. for a week: T1 to T7, time-frequency interval was 24 hours from T1 to T7. Measurement of seedlings morphological parameters viz., seedling coleoptiles and radicle lengths were measured using a millimeter ruler. Final Germination Percentage (FGP) was calculated using following formula: FGP = (Number final germinated seeds/ Total number of seed tested) x 100. Germination Energy (GE) was calculated using following formula according to Ruan et al., 2002. GE (%) = (Number of germinated seeds at 4th after sowing/ Total number of seed tested) x 100. Seed Vigor Index (SVI) was calculated according to the formula described by Abdul Baki and Anderson (1973). SVI= Radicle length (cm) × Germination (%). Relative Water Content (RWC) was calculated according to Sumithra et al., 2006. RWC % = 100 x ((FW – DW) / FW). Fresh weight was measured when the germination got ceased and dry weight was observed after drying at 70°C in hot air oven. Further germination traits such as Mean Germination Time (MT) Mean Germination Rate (MR) Coefficient variation of germination time (CVt), Synchrony (Z) was calculated according to Ranal et al., 2006, 2009.

### GREEN HOUSE BASED PHENOTYPIC TRAIT EXPERIMENT

Green house experiments were conducted in temperature controlled green house at CIMMYT, El Batan, Mexico. A completely randomized design experiment at greenhouse was conducted with 4 salinity concentrations (50mM, 100mM, 150mM, 200mM) and a control (water without salt). Soil content of the pot was about 3kg with 1:2:1 proportions of Sand: Soil: Organic matter) homogenously for all pots. Each pot was filled with gravel stones (3-5mm) from bottom up to 5cm to facilitate excess water drainage. Totally 90 pots were filled as designed experiment consisted of 6 varieties with 3 replicates for 4 concentration of salt stress with a control.

#### Pot water holding capacity (WHC) measurements

For measuring water holding capacity, 10 random pots were chosen and soil plus pot’s net weight was measured without adding water. Thereafter the pots were filled completely with excess water. The pots were then left for an hour to drain excess water and pots net weight was measured twice a day (morning and afternoon). Likewise, pots were measured continuously for 4 days. One-way ANOVA analysis was done resulted with the P Value 1.28E-26, P<0.001. Water holding capacity of pots for each day depended upon the evaporation rate and ambient temperature. This experiment was useful to select a pot with an appropriate and adequate soil without excess water and salt leakage to minimize the errors due to deviation of physico-chemical properties in a controlled ambient.

#### Preparation of seedlings

Five seeds of bread wheat lines viz., C306, HDR-77, Kharchia, OasisF86, WK1204 and a durum wheat line DWK26 were placed in Petri plate with appropriate amount of water to achieve germination. Seedlings of each line were allocated to blocks of pots as mentioned below with appropriate genotype and salt concentration labelling. After a week, thinning was done to leave a single seedling per plot. Five pots were allocated for the concentrations of 0 (control), 50mM, 100mM, 150mM and 200mM for each variety and three replicates were maintained. Therefore, in total 90 pots were completely randomized inside the greenhouse to have a homogenous ambient.

#### Phenotypic traits measurement

The phenotypic traits including Plant Height (PH) (cm), Number of Tillers (NT), Number of Effective Tillers (NoET), Flag Leaf Length (FLL), Total Number of Spikes (TS), Spike Length (SL) were measured. Booting or heading was measured 50 days after sowing (DAS) from control and salt stressed genotypes. Statistical Analysis: Analysis of variance (ANOVA) was carried out and Least Significant Difference (LSD) with Bonferroni correction was estimated using R (R Core Team, 2014).

#### Statistical Analysis

Analysis of variance (ANOVA) was carried out and Least Significant Difference (LSD) with Bonferroni correction was estimated using R (R Core Team, 2014).

## RESULTS

### *HKT1;5* gene sequence characterization

Amplification of salinity tolerance gene *TaHKT1;5-D* was done in diverse wheat lines (Fig. 2). Amplification was achieved in all 16 lines of bread wheat. Subsequently, 12 selected lines were selectively sequenced (based on preference with salinity tolerance, sensitiveness and/or with other interesting features such as drought/heat tolerance etc) comprising four alleles from bread wheat. Remaining 17 sequenced accessions were wheat wild relatives and ditelosomic lines. Eight of the sequences are ∼3 kb size from bread wheat lines, with complete CDS-ORFs. Rest of the 21 sequences are partial CDS and/or 5’ UTR of ∼1kb size from wheat or wild relatives. Gene sequence of bread wheat varieties, wild/progenitor species, durum wheat and ditelosomic lines was submitted at GenBank (Table 2).

**Table 2.** NCBI GenBank accession numbers with allele description of *TaHKT1;5* gene from diverse collection of *T. aestivum* cultivars and wild relatives.

| Wheat/wild relatives | Allele | GenBank | Size (bp) | SNPs/Allele Description |
| --- | --- | --- | --- | --- |
| *T. aestivum* WK1204 | A | KU184266 | 3073 | g.[830T>C;832T>C;2106;T>C] |
| *T. aestivum* Dharwar Dry | B | KU212871 | 2826 | g.[830T>C;832C>T;2106;T>C] |
| *T. aestivum* Munal#1 | B | KU184267 | 2438 | g.[830T>C;832C>T;2106;T>C] |
| *T. aestivum* Dharwar Dry | C | KU212872 | 2826 | g.[830T>C;832T>C;2106;C>T] |
| *T. aestivum* OasisF86 | C | KU212870 | 3068 | g.[830T>C;832T>C;2106;C>T] |
| *T. aestivum* HDR-77 | C | KU212869 | 3122 | g.[830T>C;832T>C;2106;C>T] |
| *T. aestivum* C306 | C | KU242566 | 1066 | g.[830T>C;832T>C;2106;C>T] |
| *T. aestivum* Kharchia | C | KU212875 | 3074 | g.[830T>C;832T>C;2106;C>T] |
| *T. aestivum* Sakha8 | C | KU242567 | 965 | g.[830T>C;832T>C;2106;C>T] |
| *T. aestivum* YMI#6 | C | KU242568 | 1047 | g.[830T>C;832T>C;2106;C>T] |
| *T. aestivum* Shorowaki | C | KU242569 | 965 | g.[830T>C;832T>C;2106;C>T] |
| *T. aestivum* Lu-26 | C | KU242570 | 1035 | g.[830T>C;832T>C;2106;C>T] |
| *T. aestivum* Kharchia 65 | C | KU212873 | 2883 | g.[830T>C;832T>C;2106;C>T] |
| *T. aestivum* Kharchia 65 | D | KU212874 | 2883 | g.[830C>T;832T>C;2106;C>T] |
| *A. tauschii* CWI94958SH | T1 | KU253614 | 958 | g.[724G>T] |
| *A. tauschii* CWI94956SH | T2 | KU253615 | 1009 | g.[724T>G] |
| *T.monococcum* PI272558 | M1 | KU253607 | 994 | g. [234 A>G; 745C>G; 832C>T] |
| *T. monococcum* PI272558 | M2 | KU253608 | 994 | g. [234 G>A; 745C>G; 832T>C] |
| *T. monococcum* PI428150 | M1 | KU253609 | 966 | g. [234 A>G; 745C>G; 832C>T] |
| *T. monococcum* PI326317 | M1 | KU253610 | 936 | g. [234 A>G; 745C>G; 832C>T] |
| *T. monococcum* PI264935 | M1 | KU253611 | 672 | g. [234 A>G; 745C>G; 832C>T] |
| *T. monococcum* PI272560 | M1 | KU253612 | 978 | g. [234 A>G; 745C>G; 832C>T] |
| *T. monococcum* PI272560 | M3 | KU253613 | 978 | g.[ 234 A>G; 745 G>C; 832T>C] |
| *T.monococcum* PI264935 | N1 | KY110746 | 942 | N/A |
| *T.monococcum* PI272560 | N1 | KY110747 | 893 | N/A |
| *A.speltoides* CWI94201SH | As2 | KY110745 | 632 | N/A |
| *Ditelosomic* DTGH09_1566 | DS2 | KY110743 | 902 | N/A |
| *Ditelosomic* DTGH09_1559 | DS1 | KY110744 | 879 | N/A |
| *T. durum* DWK26 | B-Td1 | KY110748 | 1007 | N/A |

**Figure 2.**
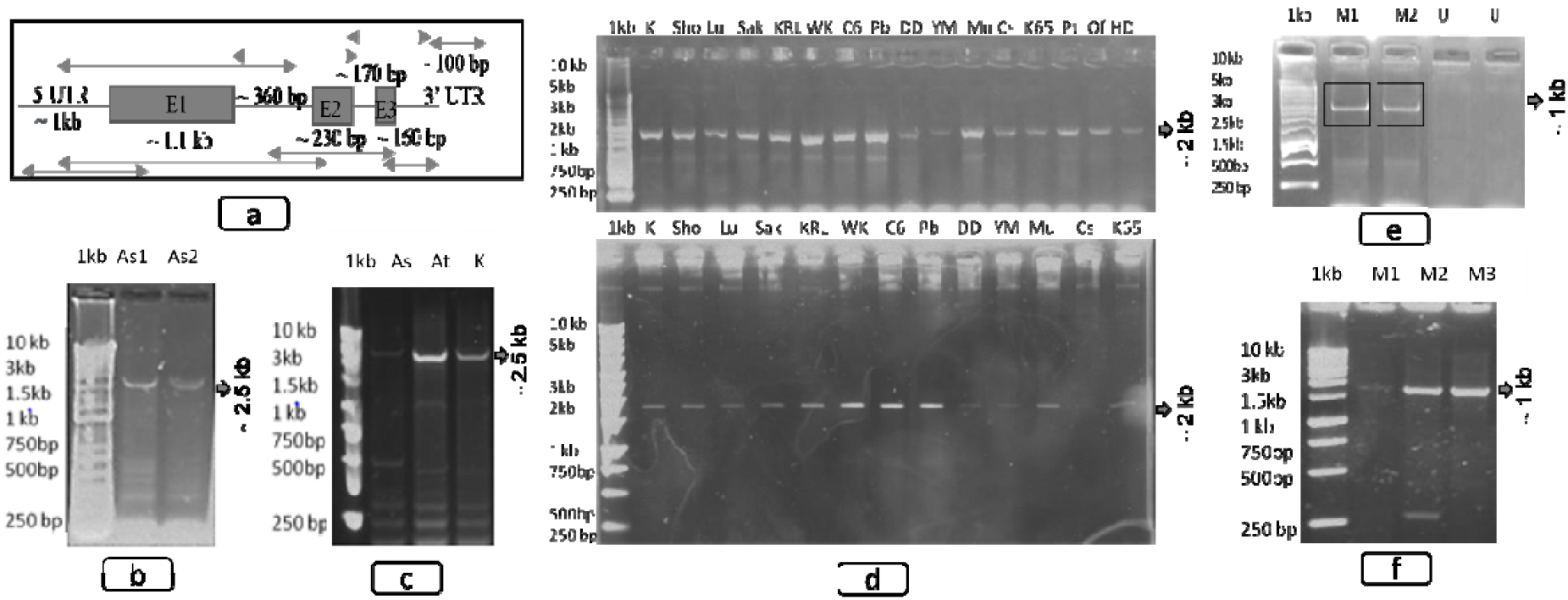
a) Schematic diagram of *HKT1;5* gene structure and approximate size of UTRs, exons, and introns. b) *HKT1;5* gene amplified in *A. speltoides*. c) *HKT1;5* gene amplified in *A. speltoides, A. tauschii* and bread wheat variety Kharchia d) Top gel: Amplification of *HKT1;5* gene in diverse *T. aestivum* varieties, K: Kharchia, Sho: Shorowaki, Lu: Lu26, Sak: Sakha, KRL: KRL1-4, WK: WK1204, C6: C306, Pb: Pasban 90, DD: Dharwar dry, YM: YMI#6, Mu: Munal#1, CS: Chinese spring, K65: Kharchia65, Pt: Pastor, Of: Oasis F86, HD: HDR-77. Bottom Gel: Purified fragments showing the intact banding pattern of the first 13 samples from top gel. e) *HKT1;5* gene amplified in *T. monococcum*, but not amplified in *T. urartu*. f) *HKT1;5* gene amplified in *T. monococcum*.

The PCR amplification derived sequencing and allele mining in bread wheat detected three SNPs, among them two SNPs at 830^th^ and 832^nd^ positions located at 5’UTR and third one located at 2106^th^ position of first intron with reference to Byrt et al., 2007. The gene is located at chromosome 4DL between 507964629 bp to 507967781 bp (TraesCS4D02G361300) that revealed with BLASTN at wheat ENSEMBL. We found four alleles were designated as A, B, C and D among 12 geographically diverse bread wheat cultivars and allele ‘C’ is prevalent (Table 1). Variant Effect Predictor revealed that 67% upstream variants (Genomic positions: 507965458 bp, 507965460 bp) and 33% intron variants (Genomic position: 507966734 bp) without amino acids change as expected for corresponding SNPs positions at 830, 832 and 2106 on the sequence of *TaHKT1;5* gene. However, a missense variant (506^th^ Serine/Glycine) with deleterious mutation (SIFT score: 0.05) was observed in exon 3 as an inter-homoeologous variant (Genomic position: 507967636) from *TaHKT1;5* gene between chromosome 4DL to 7BS (ENSEMBL: BA00457515 SNP) comparison.

Identified SNPs reflect the homozygotic nature of alleles in all the varieties with an exception in varieties Dharwar Dry (alleles ‘B’ and ‘C’) and Kharchia 65 (alleles ‘C’ and ‘D’) whereby a heterozygotic nature of the allele was observed suggesting the existence of an alternative variant of the gene *TaHKT1;5-D*. To check the allelic diversity in wild relatives, we used species-specific primers developed for *T. monococcum* and *T. aestivum* 4DL or *A. tauschii*. These primers successfully amplified *TaHKT1;5-D* gene in *T. monococcum, A. speltoides* and *A. tauschii*. The species-specific primers from *T. monococcum* amplified a native *TmHKT1;5-A* gene itself (M allele), also an ortholog in *A. tauschii* (Amplified, though not sequenced as M allele related *A*.*tauschii* (named as putative *At_HKT*) allele sequence directly retrieved from public database). Likewise, *T. aestivum* 4DL or *A. tauschii* specific primers also amplified a native *AtHKT1;5-D* gene (Allele ‘T’) and an ortholog from *T. monococcum* (N allele). These results highlight the existence of two major alleles ‘M’ and ‘N’ in *T. monococcum*. Interestingly, we could not amplify any orthologs in *T. urartu*. Based on SNPs within the M allele, three sub-alleles were detected, designated as M1, M2, M3, while major allele N exhibited only a single variant (N1) (Table 1). Also, two major alleles were identified in *A. tauschii*. The first allele was designated as ‘T’ and consists of two sub-alleles, T1 and T2 and the second allele (of *A*.*tauschii* related to M allele of *T. monococcum*) was derived from a public database IWGSC-URGI (https://wheat-urgi.versailles.inra.fr/Seq-Repository/BLAST) and named as putative *At_HKT*.

The comparison between major alleles (M and N from *T. monococcum*, T and *At_HKT* from *A. tauschii)* exhibited the polymorphism at 5% level, while the comparison within sub-category of major alleles called minor alleles showed 0 to 0.2% polymorphism. The phylogenetic tree, constructed based on maximum likelihood with 1000 bootstrap, clearly revealed some distinct clusters of major and minor alleles (Fig. 3a). In 5’ UTR region, a continuous stretch of adenine was observed at higher number in *A. tauschii* gene (*At_HKT* allele) from 554 bp onwards, whereas in *T. aestivum* and *T. monococcum* the repetition of adenine was lower. One variant from *A. tauschii* (*At_HKT*) is a major source of the 4DL chromosome bearing *HKT 1;5* gene to *T. aestivum* as it shown close phylogenetic relationship each other. (Fig. 3b). Distance matrix of these alleles from corresponding species is shown in Supplementary Table 2. The alignment around exon1 between wheat and *A. tauschii* indicates the vicinity of evolutionary relatedness between At_HKT allele and alleles of wheat, while T allele (T1, T2) lies farther (Fig. 3c). Relative divergence time concerning *HKT1;5* gene is recent for Triticeae members in Pooideae subfamily in Poaceae than model grass *Brachypodiuum distachyon* and other species belongs to the subfamily of Oryzoideae and Panicoideae as shown in Fig. 3d.

**Figure 3.**
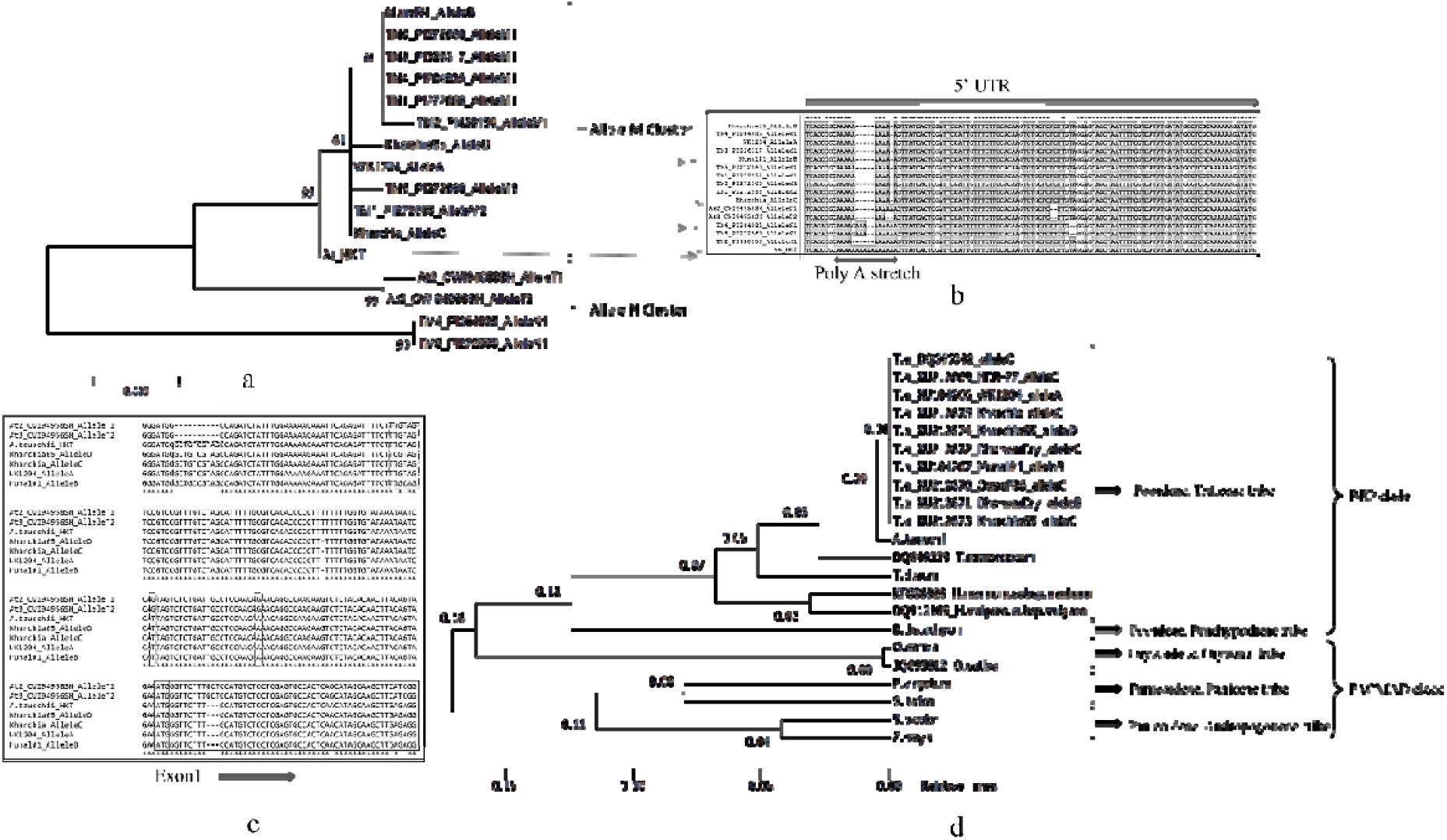
a) Phylogenetic comparison of M and N alleles from *T. monococcum* to *T. aestivum* and *A. tauschii* alleles. b) Alignment showing Poly A stretches in 5’ UTR and nucleotide comparison among the species/sequences. c) Interspecific SNPs and insertion/deletion mutations between *A. tauschii* and *T. aestivum* alleles. Large box indicates exon1 start codon and an insertion. Dotted box indicates the difference between *A. tauschii* alleles T1, T2 and *A. tauschii* allele ‘*At_HKT*’ sequence. Shaded box indicates SNPs difference from *T. aestivum*. d) *HKT1;5* gene based Relative Divergence time among specific tribes in Poaceae.

Concerning *A. speltoides*, amplification was achieved in some accessions accession, it perhaps due to accumulated mutations in and around primer binding sites. The *HKT1;5* gene partial sequence generated from *A. speltoides* (Allele ‘As2’) revealed an alteration of amino acids in primary structure of protein due to the insertion of four nucleotides (CGCG) caused a frameshift mutation at exon1 suggesting a distinct allelic variant (Genbank: KY110745, Fig. 4a). Besides, a BLASTN search for additional alleles with top score sequences from IWGSC-URGI (https://urgi.versailles.inra.fr/Species/Wheat), revealed the existence of four more alleles in *A. speltoides* with available genomic resource. Hence, there is a need of more sequencing effort in natural accessions/populations of this species to excavate further alleles. None of the database sequences showed a frameshift mutation with CGCG insertion at exon1 as ‘As2’ allele of *A. speltoides*. The BLASTN of the sequences (https://urgi.versailles.inra.fr) revealed that the complete gene sequence was present only on homeolog chromosome 4D. To better understand the number of gene variants on chromosome 4D, we used two ditelosomic lines, the first one called DTGH09_1559 possessed 4D chromosome long arm (L) but lacking 4D chromosome short arm (S) (Allele ‘DS1’) and the second one namely DTGH09_1556 (Allele ‘DS2’) was the opposite of DTGH09_1559 i.e. possessing 4D chromosome short arm (S) but lacking 4D chromosome long arm (L). The alignment of sequenced homeolog from DTGH09_1559 with 4DL (−4DS) (GenBank: KY110744) revealed a 99.89% identity with *T. aestivum* (GenBank: KU184266), while it was 93.25% identical to *T. durum* (GenBank: KY110748). In contrast, the homeolog from DTGH09_1556 4DS (−4DL) (GenBank: KY110743) revealed a 98.89% identity to *T. durum* while it was 93.08% identical to *T. aestivum*. Both ditelosomic lines DTGH09_1559 and DTGH09_1556 have 93.78 % identity to each other. Protein sequence alignment among bread wheat, durum wheat and two ditelosomic lines corresponding to exon1 is shown in Fig. 4b, the nucleotide alignment with an insertion and deletion is shown in Fig. 4c. This indicated that the homeolog from DTGH09_1556 4DS (−4DL) has similarity with an ortholog of durum wheat and might have amplified from ‘A’ or ‘B’ genome rather than from the D genome of ditelosomic line DTGH09_1556 4DS (−4DL). BLASTN has further confirmed that DTGH09_1556 4DS (−4DL) gene sequence was located on chromosome 4B long arm (4BL) as it showed 100% sequence identity with the scaffold TGACv1_scaffold_320406_4BL from bread wheat. The allele DTGH09_1556 (Allele ‘DS2’) resembles an existing HKT8-B2 homeolog (DQ646341) in wheat, though there are 9 SNPs spanning the gene region of 1kb between these two forms, thus it is a novel variant of B2 homeolog. Further, our phylogenetic analysis suggests that there could be 5 or more paralogs within 4B chromosome homeolog, distinct orthologs in putative B genome donor *A. speltoides* and related to B-Td1 of *T. durum*. However, the allele ‘DS1’ from DTGH09_1559 is completely identical as *HKT1;5-D* at 4DL, thus it is completely belonging to the A, B, C, D alleles.

**Figure 4:**
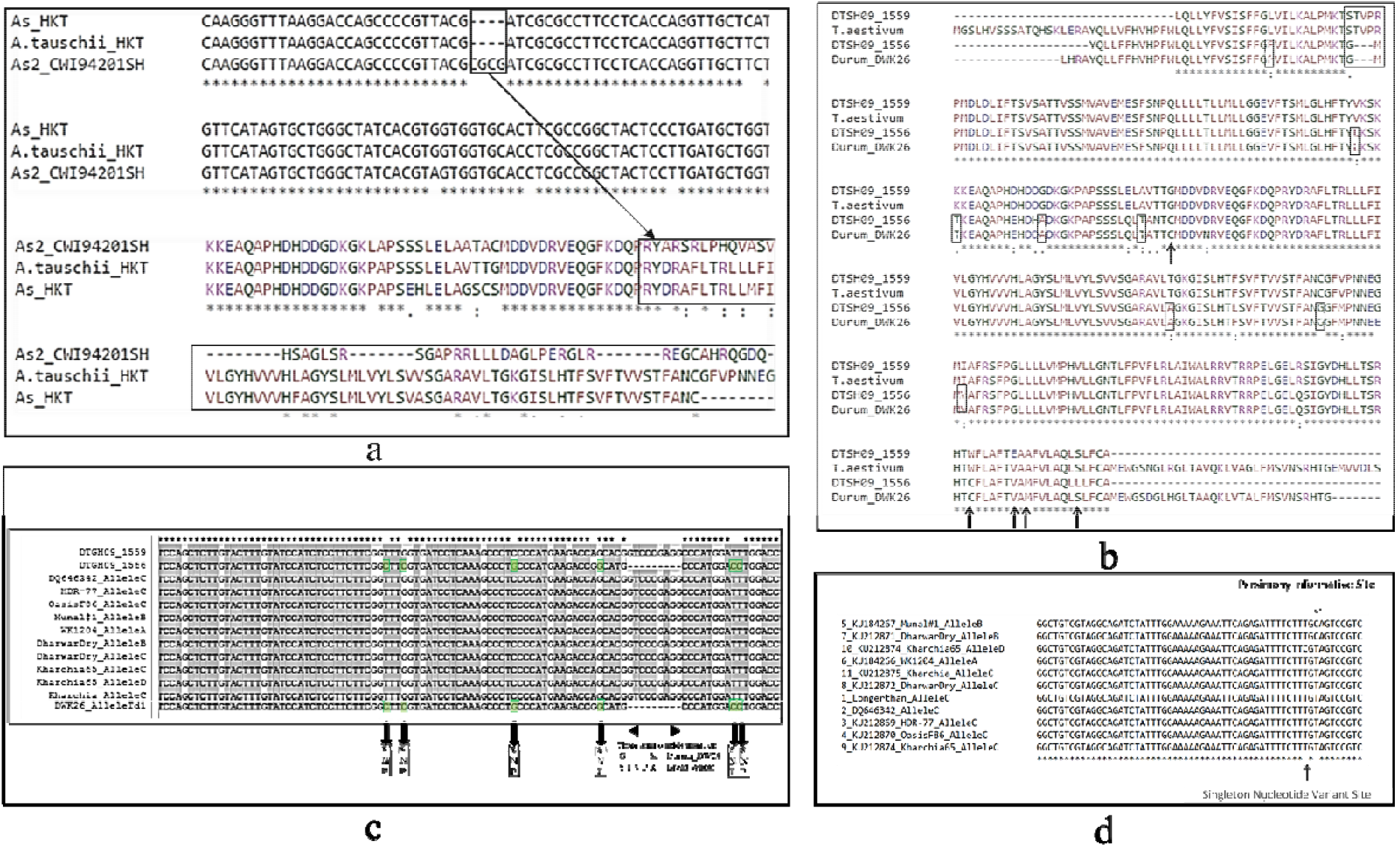
a) Interspecific SNPs insertion deletion variation between *A. tauschii* and *A. speltoides*. Box indicating the 4bp insertion caused frameshift with altered protein. b) Amino acids alignment corresponding to the region of exon1 of *HKT1;5* gene among bread wheat, durum wheat and diltelosomic lines. Small box indicates conservative amino acid changes. Large box indicates insertion deletions. Arrow indicates non conservative amino acid changes. c) Nucleotide alignment of *HKT 1;5* gene exon1 part among *T. aestivum, T. durum* and diltelosomic lines. d) Parsimony informative site and singleton nucleotide variant site in *T. aestivum* alleles.

Overall, the coding sequences of *TaHKT1;5-D* against the survey sequences of all bread wheat revealed the presence of an entire genic region on 4DL, while remnants or portions of gene regions on 4BL, 4BS and 4DS chromosomes. A non-autonomous mariner transposon element was found in intron 1 of the gene on all A, B, and D chromosomes with >3120 BLASTN hits. These results clearly indicate the mobile nature of the gene through transposition events either as a whole or part of the gene, hence multiple variants of *HKT1;5* gene in bread wheat and its progenitors is highly plausible. The presence of huge variation in sequence alignment between *T. durum* cultivar DWK26 and *T. aestivum* indicated variation between the amplified gene from 4BL and 4DL respectively. The amino acid alignment evidencing that *HKT1;5* gene sequence of durum wheat was highly comparable with diltelosomic line 1556 (+4DS, - 4DL), while *T. aestivum* was highly comparable with DTGH09_1559 (+4DS, -4DL). In *A. tauschii* the presence of two variants (*At_HKT* and T allele) revealed that these sequences sharing proximity among ditelosomic line 1559 (+ 4DL and - 4DS), bread wheat and *A. tauschii*, while Ditelosomic line 1556 (+4DS and -4DL) was differing with its own SNPs from others. The trend suggested that two paralogous variants existed in *A. tauschii* and ortholgous to wheat. Besides, ditelosomic bread wheat lines showed the existence of 2 paralogous/homoeologous variants, which revealed that one was amplified from DT line 1559 (+4DL-4DS) while another from 1556 (+ 4DS, -4DL) of 4BL. Both the alleles from *A. tauschii* (*At_HKT* and T allele) had proximity with the sequence of DT line 1559 rather than DT1556.

### Genetic diversity analysis of the *HKT1;5* alleles in bread wheat

There were 2301 invariant sites, and 3 variant sites detected. Among the variant sites, 2 were singleton nucleotide variant sites (at position 830 in 5’ UTR and at 2106 in intron1) and 1 parsimony informative site (at position 832 of 5’ UTR) with two variants. The variant sites at 5’ UTR are shown in Fig. 4d. Diversity analysis showed low polymorphism and recombination rate for this gene among the bread wheat lines concerning these four alleles in *TaHKT1;5-D* (Table 3). Gene flow and genetic differentiation showed similar trend with least effective migrants (0.13) F_ST_ (0.65) among the four alleles of the defined population/genotypes. It further elucidates the least frequency of recombination in this gene owing to autogamous nature and lesser allelic variation within bread wheat lines. The conservative nature of the sequences with lower number of mutations had resulted low haplotype diversity of the four alleles. The observed total nucleotide difference was also significantly lower (Table 4).

**Table 3.** Polymorphic sites and DNA polymorphism. h-Number of haplotypes, hd-Haplotype diversity, SD-Standard deviation, θ-4Nµ, where N is effective population size and µ is mutation rate per site per generation, π-Nucleotide diversity, η –Total number of mutations, S-Segregating sites, R-Recombination, NR-No Recombination, k-Average number of nucleotide differences, Sq-Sequence.

| H | Hd | Variance | SD | $\pi$ | $\theta$ from $\eta$ |
| --- | --- | --- | --- | --- | --- |
| 4 | 0.694 | 0.02162 | 0.147 | 0.00036 | 0.00048 |
| $\theta$ from S | Variance $\theta$ (NR) | SD $\theta$ (NR) | Variance $\theta$ (R) | SD $\theta$ (R) | $\theta$ from $\pi$ |
| 0.00048 | 0.0000001 | 0.00032 | 0.0000001 | 0.00028 | 0.00036 |
| $\theta$ from S | $\theta$ from $\eta$ | k | $\theta$ from S/Sq | Variance $\theta$ (NR)/Sq | Variance $\theta$ (R)/Sq |
| 0.00048 | 0.00048 | 0.833 | 1.104 | 0.545 | 0.406 |

**Table 4.**
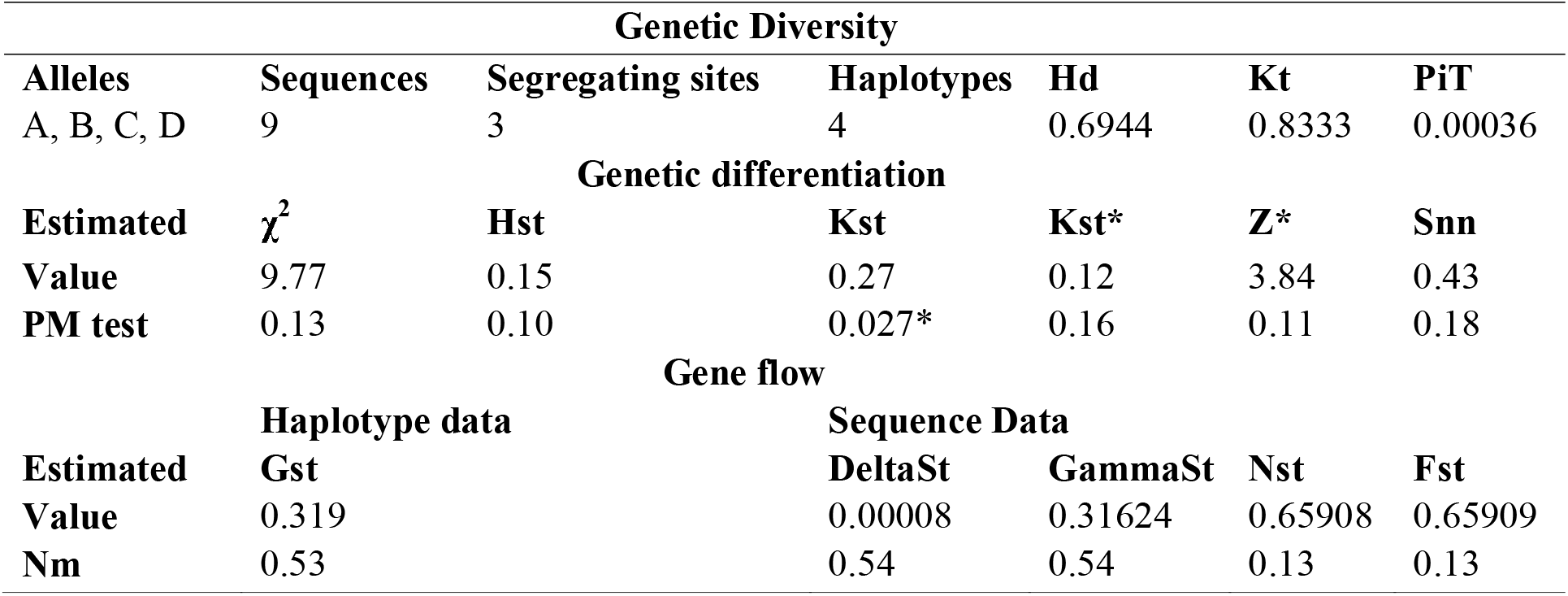
Genetic diversity, differentiation and gene flow analysis from B and C Alleles of *TaHKT1;5-D* gene. Hd-Haplotype Diversity, Kt-Average Nucleotide Differences, PiT-Nucleotide diversity, χ^2^-Chi Square test, Hst-Haplotype based statistics, Kst-Sequence based statistics, Z*-Rank Statistics, Snn-Near Neighbour Statistics, Gst-Differentiation of population, GammaSt-Gamma Statistics, Nst-N statistics, Fst-Fixation index, Nm-Effective number of migrants, PM-Permutation test with 1000 replicates, * 0.05<P.

The percentage of nucleotides was 22.96% (A), 27.73% (T/U), 26.83% (C), and 22.48% (G) derived from the alleles corresponding to 2304 positions. The C to T transition mutation (pyrimidines) substitution pattern was higher (50.59%) than other type of transition mutations. Besides, the transversion mutations (purine to pyrimidine or vice versa) were lower between the ranges of 0.04 to 0.05 %. Among the bread wheat cultivars, SNPs were not observed in coding region of the gene, hence Ks/Ka (synonymous/non-synonymous mutation) analysis was not performed. Tajima D test (D: -0.93613) with negative value indicates purifying selection due to low frequency of variations along this gene among the bread wheat cultivars, which previse population size expansion. However, Tajima D test was not significant and thus increasing sample size may provide a better clue.

### Phylogenetic analysis of the *HKT 1;5* gene

The phylogenetic tree based on all generated gene sequences and additional Poaceae member sequences (especially tribe Triticeae) from public databases revealed the presence of five orthologous groups with each group, consisting of closely related species or from closely related chromosomes in case of polyploid species (Fig. 5). The ditelosomomic line DTGH09_1559 clustered together with the *T. aestivum* and *A. tauschii* cultivars. The alleles from *T. monococcum* and *A. speltoides* deviated from the orthologous group I and were designated as an orthologous group II. Alleles from ortholog group III existed with III sub-groups of bread wheat and durum wheat samples and have proceeded from A, B and D putative genome donors from ortholog group II. The four alleles of *A. speltoides* retrieved database comes under three ortholog groups with other related species. The orthologous group IV consisted of alleles from *A. sharonensis, A. speltoides, Hordeum spp* and *B. distachyon*. This group appeared as a distinct group from the ortholog groups I-III. Group V consists of other Poaceae members used in this study apart from Triticeae and Brachypodieae tribes.

**Figure 5.**
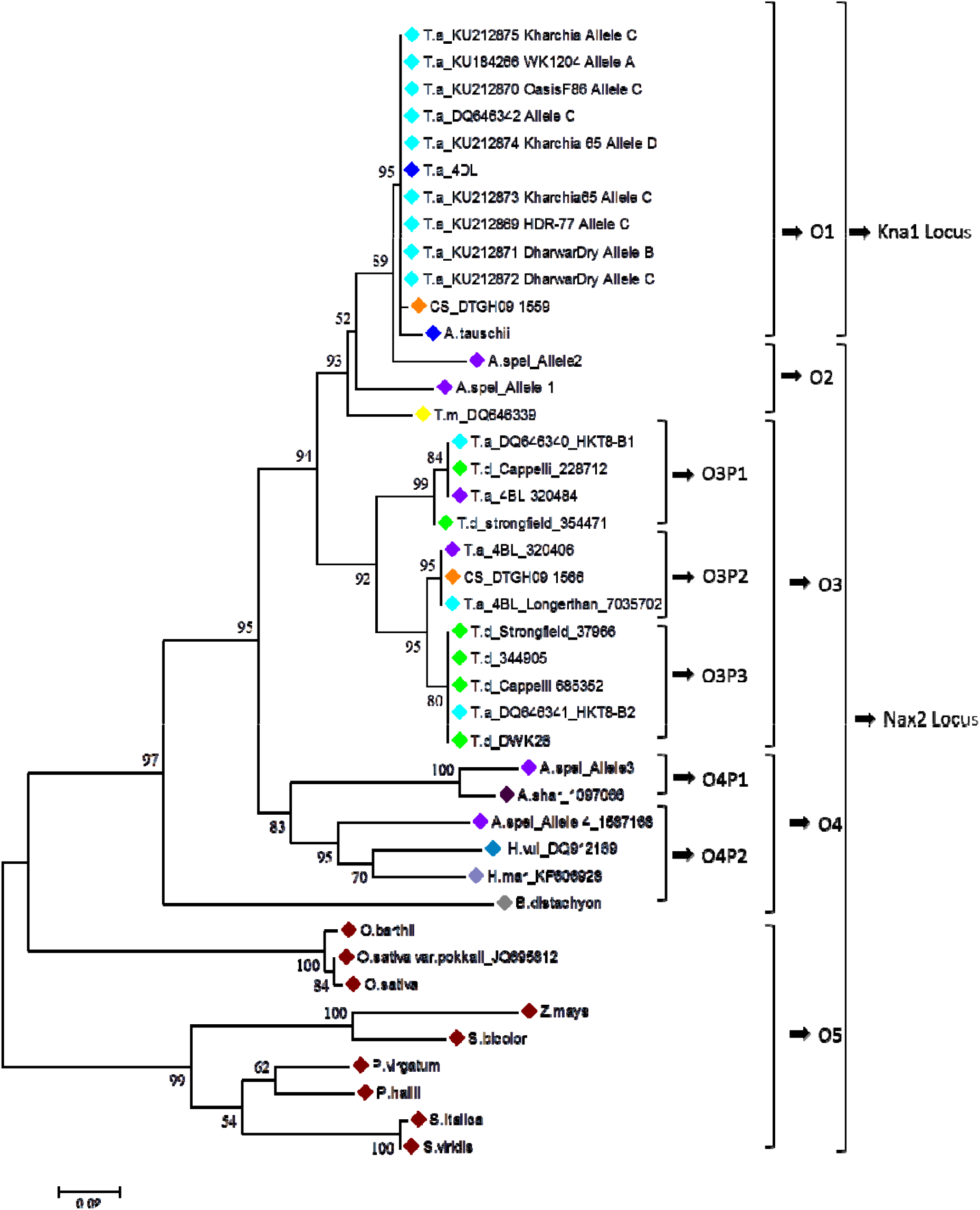
Reconstructed phylogenetic tree showing the orthologous and paralogous comparison of *HKT1;5* gene with *Kna1* and *Nax2* homoeologs relevancy in different species. O=Ortholog, P=Paralog. Phylogeny based on Maximum Likelihood method with Jukes Cantor (JC) model and 1000 bootstrap resampling.

### Comparative analysis between *T. aestivum* and *A. thaliana*

Pairwise alignment between *T. aestivum HKT1;5-D* and *A. thaliana HKT1* gene and its protein revealed the similarity at the level of 54.9 % and 48.4% respectively exonic region and corresponding protein sequence of these orthologs. Since it was intriguing, the protein modeling and comparative study was done, and it indicates the structural conservation of protein sequence between *A. thaliana* and *T. aestivum* (Fig. 6a).

**Figure 6.**
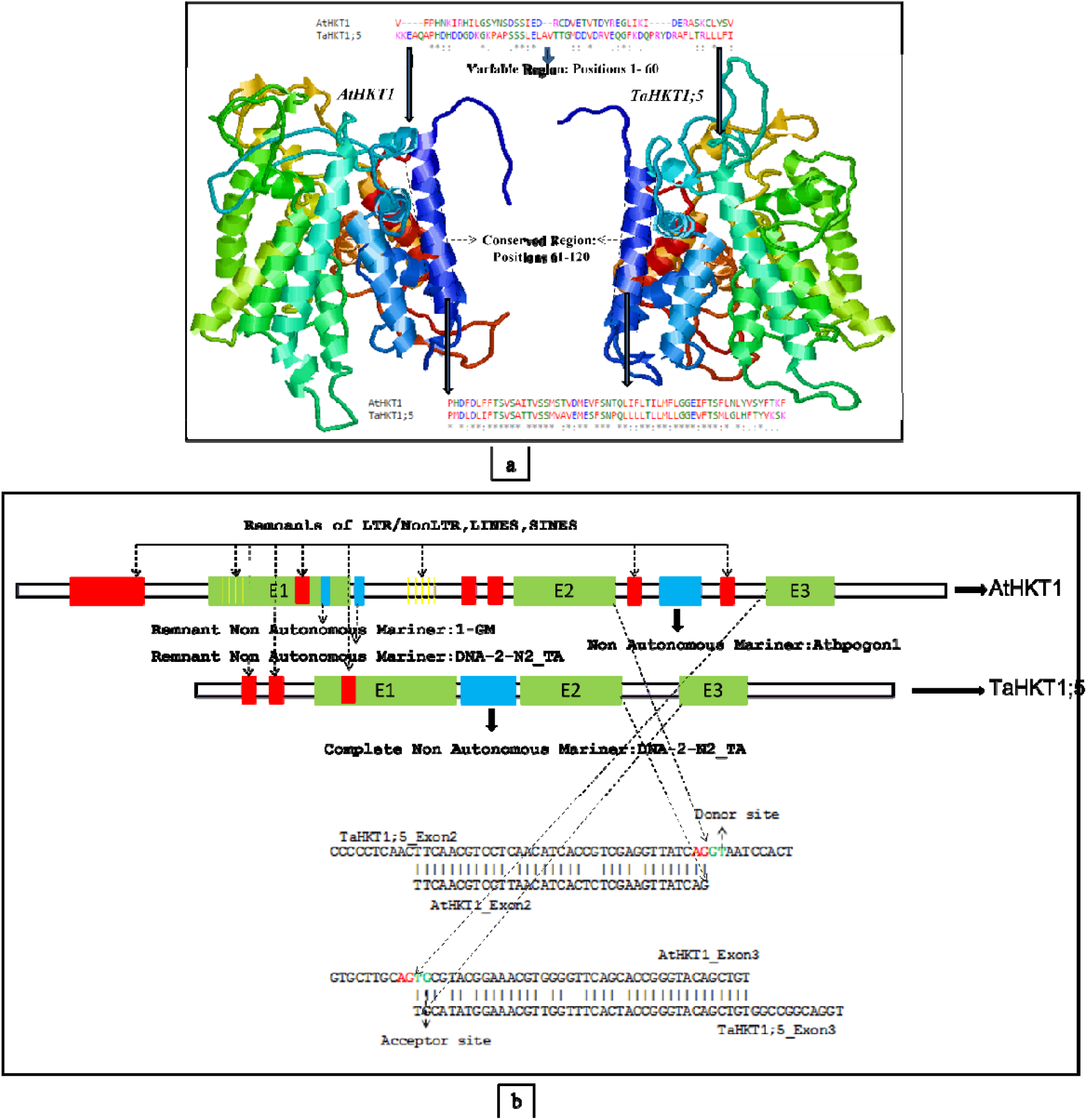
a) Protein homology modelling, tertiary, and primary structures comparison of *HKT1* protein from *A. thaliana* with *HKT1;5* from *T. aestivum*. b) Structural organization and insertions of transposons/fragments of transposon in *HKT1* gene from A. *thaliana* and *HKT1;5* from *T. aestivum*.

Nucleotide level global alignment with LAGAN revealed the presence of higher conservation in coding or exonic part of *HKT1* sequences between wheat and *A. thaliana*. The comparison of transposable elements inside the gene showed that *A. thaliana HKT1* gene has higher number of transposon insertions or remnants of transposons inside the gene compare to bread wheat *HKT1;5* gene. The conservation of CDS region, donor and acceptor splicing sites proved the similarity of the structural organization of this gene between *A. thaliana* and *T. aestivum*. Introns are enriched with the presence of more remnants of different transposon; however, a complete non-autonomous transposon called Mariner at intron1 in bread wheat was comparable between these two-diverse species (Fig. 6b). Physico-chemical properties of the protein comparison between *A. thaliana* and *T. aestivum* revealed that higher similarity of this orthologous gene from these two-diverse species (Supplementary Table 3). The structural organization of the corresponding putative protein sequences is also in similar trend. The instability index (II) indicates that *HKT1* protein from *T. aestivum* and *A. thaliana* are stable.

### Salinity stress induced phenotypic evaluation in selected allele designated wheat lines

To understand the effect of salinity during germination at seedling stage, the following experimental test was done using drought/heat tolerant lines (C306, HDR-77, so far unknow for salinity tolerance or sensitiveness), WK1204 (a yellow rust resistant line, so far unknown for salinity tolerance or sensitiveness) and a salinity sensitive line OasisF86. This test was done to understand the level of salinity tolerance in unknown varieties and these lines were further validated with subsequent experimental tests with known salinity tolerant lines such as Kharchia, Kharchia 65 etc.

Detailed description of the of the trait variability with LSD in the HDR-77 (Allele ‘C’), C306 (Allele ‘C’), Oasis F86 (Allele ‘C’) and WK1204 (Allele ‘A’) *T. aestivum* cultivars is given in Table 5. The germination test among these four cultivars showed OasisF86 (‘C’ allele) and WK1204 (‘A’ allele) to be early germinating than other two cultivars with least germination Mean Time (MT). Final Germination Percentage (FGP) was maximum in C306 and HDR-77, even at 200mM salt stress, suggesting that the seedlings of these two varieties are more tolerant to salt stress than others. Fresh Weight (FW) augmented upon increasing the salinity concentration, maximum at 50-100 mM level than control in all varieties, similarly FW was increased with increasing salt concentration, reached maximum at 200mM for all especially for HDR-77. (Fig. 7).

**Table 5.** Mean values and variability of traits under different treatments of salinity stress including control and mean values and variability of each genotype with underlying phenotypic features responding to treatment. Letters not sharing similarity are statistically significant to LSD in each row at P=0.05 level. FGP= Final Germination Percentage, GE= Germination Energy, SVI= Seed Vigor Index, CL= Coleoptile Length, RL= Radicle Length, NoR= Number of radicles, ColL-Leaf Length from Coleoptile, FW= Fresh Weight, DW= Dry Weight, RWC= Relative Water Content, MT= Mean Germination Time, CVt= Coefficient Variation of Germination Time, MR=Mean Germination Rate, U= Uncertainty of the Germination Process, Z=Synchrony. Statistical significance: *P<0.05, **p<0.01, ***P<0.001.

| Traits | Treatment |  |  |  |  | LSD at P=0.05 | F test significance |
| --- | --- | --- | --- | --- | --- | --- | --- |
|  | Control | 50 mM | 100 mM | 150 mM | 200 mM |  |  |
| **FGP** | 96.11a | 88.88ab | 88.33ab | 86.66b | 83.88b | 7.8 | 0.03957 * |
| **GE** | 13.33a | 11.11a | 10.55a | 8.88a | 5.55a | 8.68 | 0.47182 |
| **SVI** | 998.66a | 891.99a | 635.48b | 396.34c | 255.15d | 116.95 | < 2e-16 *** |
| **CL** | 44.68a | 43.72ab | 38.61bc | 37.68c | 36.95c | 5.72 | 0.0205 * |
| **RL** | 103.60a | 100.14a | 72.20b | 44.45c | 29.67d | 12.36 | < 2e-16 *** |
| **NoR** | 3.98a | 3.98a | 3.79ab | 3.75ab | 3.59b | 0.26 | 9.31e-07 *** |
| **Coll** | 104.49a | 99.40a | 82.81b | 70.86c | 42.47d | 8.94 | 4.39e-16 *** |
| **FW** | 1.47a | 1.42a | 1.34a | 1.33a | 1.08b | 0.15 | 8.83e-05 *** |
| **DW** | 0.50a | 0.49a | 0.49a | 0.43b | 0.42b | 0.04 | 0.000918 *** |
| **RWC** | 68.93a | 66.27ab | 63.00bc | 61.91bc | 59.10c | 5.38 | 0.00613 ** |
| **MT** | 3.24a | 3.02a | 2.68b | 2.67b | 2.65b | 0.25 | 2.51e-05 *** |
| **CVt** | 7.71a | 7.11a | 7.11a | 6.41a | 6.39a | 1.66 | 0.4682 |
| **MR** | 0.38a | 0.38a | 0.38a | 0.34b | 0.31b | 0.03 | 0.00052 *** |
| **U** | 1.70a | 1.62a | 1.58a | 1.53a | 1.52a | 0.32 | 0.824 |
| **Z** | 29.97a | 25.50ab | 22.77ab | 19.20b | 15.94b | 10.31 | 10.0772 |
| **Genotypes** |  |  |  |  |  |  |  |
| **Traits** | **C306** | **HDR-77** | **WK1204** | **Oasis F86** | **LSD at P=0.05** | **F test Significance** |  |
| **FGP** | 96.88a | 90.22ab | 84b | 84b | 6.98 | 0.00108 ** |  |
| **GE** | 18.22a | 9.77b | 8.00b | 3.55b | 7.76 | 0.00393 ** |  |
| **SVI** | 820.48a | 717.02a | 520.01b | 484.58b | 104.6 | 2.47e-08 *** |  |
| **CL** | 49.14a | 44.55a | 34.06b | 33.55b | 5.11 | 3.15e-08 *** |  |
| **RL** | 84.52a | 79.01a | 60.44b | 56.08b | 11.05 | 2.75e-06 *** |  |
| **NoR** | 4.23a | 3.89b | 3.67bc | 3.50c | 0.23 | 9.31e-07 *** |  |
| **ColL** | 94.15a | 90.73a | 77.30b | 65.85c | 7.99 | 4.99e-09 *** |  |
| **FW** | 1.44a | 1.41a | 1.36a | 1.10b | 0.14 | 2.49e-05 *** |  |
| **DW** | 0.55a | 0.50b | 0.48b | 0.34c | 0.04 | 1.73e-13 *** |  |
| **RWC** | 68.24a | 65.47ab | 62.08bc | 59.59c | 4.81 | 0.00416 ** |  |
| **MT** | 3.57a | 2.78b | 2.64bc | 2.43c | 0.22 | 6.04e-13 *** |  |
| **CVt** | 10.12a | 7.52b | 5.08c | 5.07c | 1.49 | 5.95e-09 *** |  |
| **MR** | 0.41a | 0.38ab | 0.36b | 0.28c | 0.03 | 4.21e-10 *** |  |
| **U** | 1.93a | 1.75a | 1.38b | 1.29b | 0.29 | 8.45e-05 *** |  |
| **Z** | 36.33a | 30.44a | 16.24b | 7.70b | 9.22 | 1.76e-07 *** |  |

**Figure 7.**
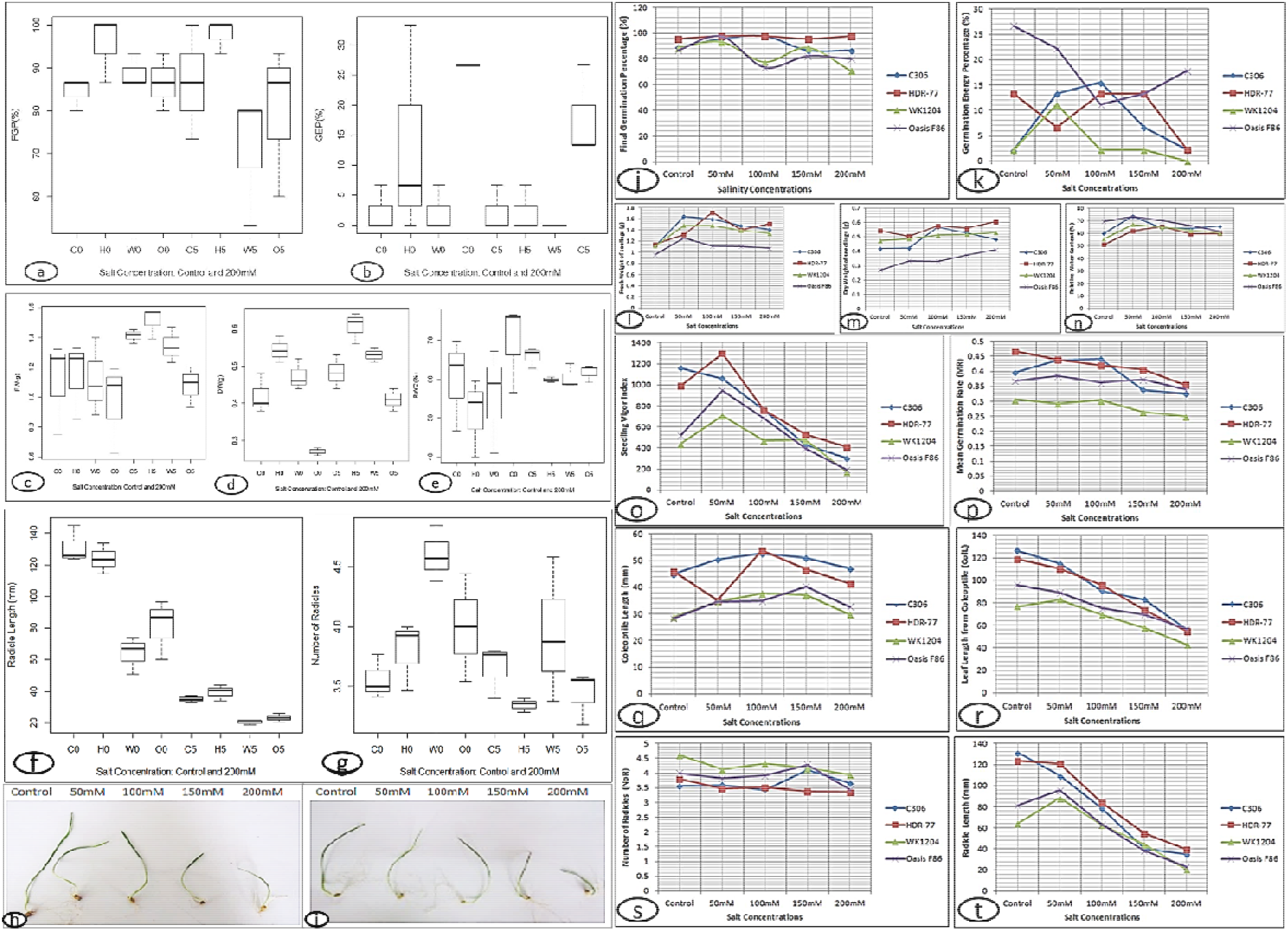
a-g. Box plot showing the distribution of variation for traits FGP, GEP, FW, DW, RWC, RL and NoR among the cultivars between control and high stress (200 mM). Legend: C0-C306 Control, H0-HDR-77 control, W0-WK1204 control, O0-Oasis F86 control; C5-C306 200 mM, H5-HDR-77 200 mM, W5-WK1204 200 mM, O5-Oasis F86 200 mM. Reduction of coleoptile size and radical size and numbers upon increasing salt stress in bread wheat HDR-77 (h) and C306 (i). j-t. Measured traits plotted against salinity concentration from 0 to 200mM for cultivars: C306, HDR-77, WK1204, OasisF86.

At 200 mM salinity stress, all the varieties exhibited higher dry (DW) weight at 200 mM than control; notably HDR-77 exhibited the highest DW than the rest of the varieties. Relative Water Content (RWC) slightly decreased by increasing salt concentration; and, at 200 mM salinity stress. RWC was higher for C306 than rest of the varieties. RWC was comparatively higher at control and lower at 200mM for OasisF86, it suggests that the salinity sensitive variety Oasis F86 was able to absorb and retain the water efficiently than rest of the cultivar in the absence of salinity stress. Overall RWC was higher in the tolerant lines (C306 and HDR-77). Radicle Length (RL) varied or decreased corresponding to increased salinity concentration, proving the inversely proportional relationship between these two factors. Similar trend was observed with Number of Radicles (NoR). In absence of salinity stress (control) RL was longer in C306 and HDR-77 than WK1206 and OasisF86; while in contrast at control, the NoR were less in C306 and HDR-77 than in WK1206 and OasisF86. It clearly suggests that at control, the salt tolerant lines were able to produce longer radicle (RL) than sensitive lines, while sensitive lines produce more radicles (NoR) than salt tolerant lines. This finding suggests that the sensitive line WK104 with allele ‘A’ is highly comparable for its sensitivity with specific feature of producing more radicles (NoR) under salinity stress than lines with Allele ‘C’. (Fig. 7).

Mean Germination Time (MT) and Mean Germination Rate (MR) for both sensitive cultivars (OasisF86 and WK1206) were shorter than other two varieties. It suggests that sensitive lines tend to germinate faster or earlier and are more susceptible to salt stress than HDR-77 and C306. This analysis revealed that the cultivar C306 is most tolerant to salinity, C306 is in the second most tolerant cultivar. The allele ‘C’, was common to both salinity tolerant and susceptible cultivars such as Kharchia and OasisF86, while only one genotype WK1204 had allele ‘A’, which is saline sensitive cultivar as revealed with our analysis. Dharwar Dry is a salinity sensitive cultivar possessed both ‘B’ and ‘C’ alleles. Similarly, the salinity tolerant cultivar Kharchia 65 possessed both ‘C’ and ‘D’ alleles. It is intriguing that cultivars showed sensitiveness had an allele ‘A’ and some of them with allele ‘C’, while the combination of alleles ‘B’ (g.[830T>C;832C>T;2106;T>C]) or ‘D’ (g.[830C>T;832T>C;2106;C>T]) present in *T. aestivum* cultivar Kharchia 65), which is a known saline tolerant cultivar. (Fig. 7). Further similar experimental tests derived result among the wheat varieties Kharchia, Kharchia65, OasisF86, DharwarDry and DWK26 is shown in Supplementary Table 4, Supplementary Table 5, Supplementary Figure1.

Greenhouse based phenotypic analysis was carried out for the bread wheat cultivars and a durum wheat and results reported. Statistical results concerning wheat cultivars to salinity stress is listed in table 5. Mostly saline sensitive bread wheat cultivars OasisF86 (Allele ‘C’), WK1204 (Allele ‘A’) and durum wheat cultivar DWK26 was observed with earlier booting or heading stage in short period (50 DAS) of the time since the salinity exposure. However, salinity stress didn’t induce early (50 DAS) booting and heading in salinity tolerant variety bread wheat variety Kharchia (Allele ‘C’), drought tolerant and heat tolerant varieties HDR-77 (Allele ‘C’) and C306 (Allele ‘C’) (Supplementary Table 6). The greenhouse-based experiment also revealed that the saline sensitive cultivars tend to have an early maturation and short life cycle revealed through the observation and measurement of booting and heading at 50 DAS (Days After Sowing). In addition, in accordance with salt tolerant reference line Kharchia, the drought and heat tolerant cultivars were exhibited better salinity tolerance. Even the level of tolerance for salinity was slightly higher for the lines C306 and HDR-77 than Kharchia (Table 6).

**Table 6.** Mean values and variability of traits under different treatments of salinity stress including control and mean values and variability of each genotype with underlying phenotypic features responding to treatment. Letters not sharing similarity are statistically significant to LSD in each row at P=0.05 level. PH=Plant Height, NoT=Number of Tillers, NoET=Number of Effective Tillers, FLL=Flag Leaf Length, TS= Total Number of Spikes, SL= Spike Length. Statistical significance: .p<1, *P<0.05, **p<0.01, ***P<0.001, Ns-Not significant.

Treatment
| Traits | Control | 50 mM | 100 mM | 150 mM | 200 mM | LSD at P=0.05 | F test significance |
| --- | --- | --- | --- | --- | --- | --- | --- |
| PH | 59.44a | 59.27a | 57.72a | 56.16a | 47.88b | 7.08 | 3.20e-05 *** |
| NoT | 5.05a | 4.72a | 4.66a | 4.50a | 4.38a | 1.15 | 0.52 Ns |
| NoET | 4.11a | 3.38ab | 3.38ab | 3.00b | 2.72b | 0.86 | 0.000274 *** |
| FLL | 55.19a | 50.86ab | 43.72ab | 42.58b | 41.94b | 12.57 | 0 0.00907 ** |
| TS | 4.11a | 3.38ab | 3.38ab | 3.00b | 2.72b | 0.86 | 0.000274 *** |
| SL | 25.66a | 23.22ab | 21.08ab | 19.16bc | 15.86c | 4.88 | 1.58e-06 *** |

| Genotypes |  |  |  |  |  |  |  |  |
| --- | --- | --- | --- | --- | --- | --- | --- | --- |
| Traits | C306 | HDR-77 | Kharchia | OasisF86 | WK1204 | DWK26 | LSD at P=0.05 | F test Significance |
| PH | 62.76a | 61.56a | 60.93a | 51.70b | 50.80b | 48.83b | 8.13 | 5.11e-08 *** |
| NoT | 6.60 a | 5.00 b | 4.86b | 4.53bc | 3.73bc | 3.26c | 1.33 | 9.8e-10 *** |
| NoET | 3.80a | 3.80a | 3.40ab | 3.33ab | 2.86ab | 2.73b | 0.99 | 0.004342 ** |
| FLL | 63.80a | 62.90a | 59.90a | 35.06b | 33.21b | 26.30b | 14.44 | 3.08e-15 *** |
| TS | 3.80a | 3.80a | 3.40ab | 3.33ab | 2.86ab | 2.73b | 0.99 | 0.004342 ** |
| SL | 25.43a | 22.63ab | 22.53ab | 21.36ab | 19.66bc | 14.36c | 5.6 | 2.74e-06 *** |

Statistical result from greenhouse experiment revealed that bread wheat variety C306 (Allele ‘C’) is more saline tolerant (PH: 62.76a, LSD at P=0.05, F test significance 5.11e-08, P<0.001) (Table 5) than other bread wheat varieties. Oasis F86 (Allele ‘C’) is saline sensitive cultivar and comparable with WK1204 (Allele ‘A’) and durum for salt sensitiveness (Fig. 8). Overall experiment revealed the salinity sensitiveness or tolerance among the varieties through the possible comparison of phenotypes with alleles.

**Figure 8.**
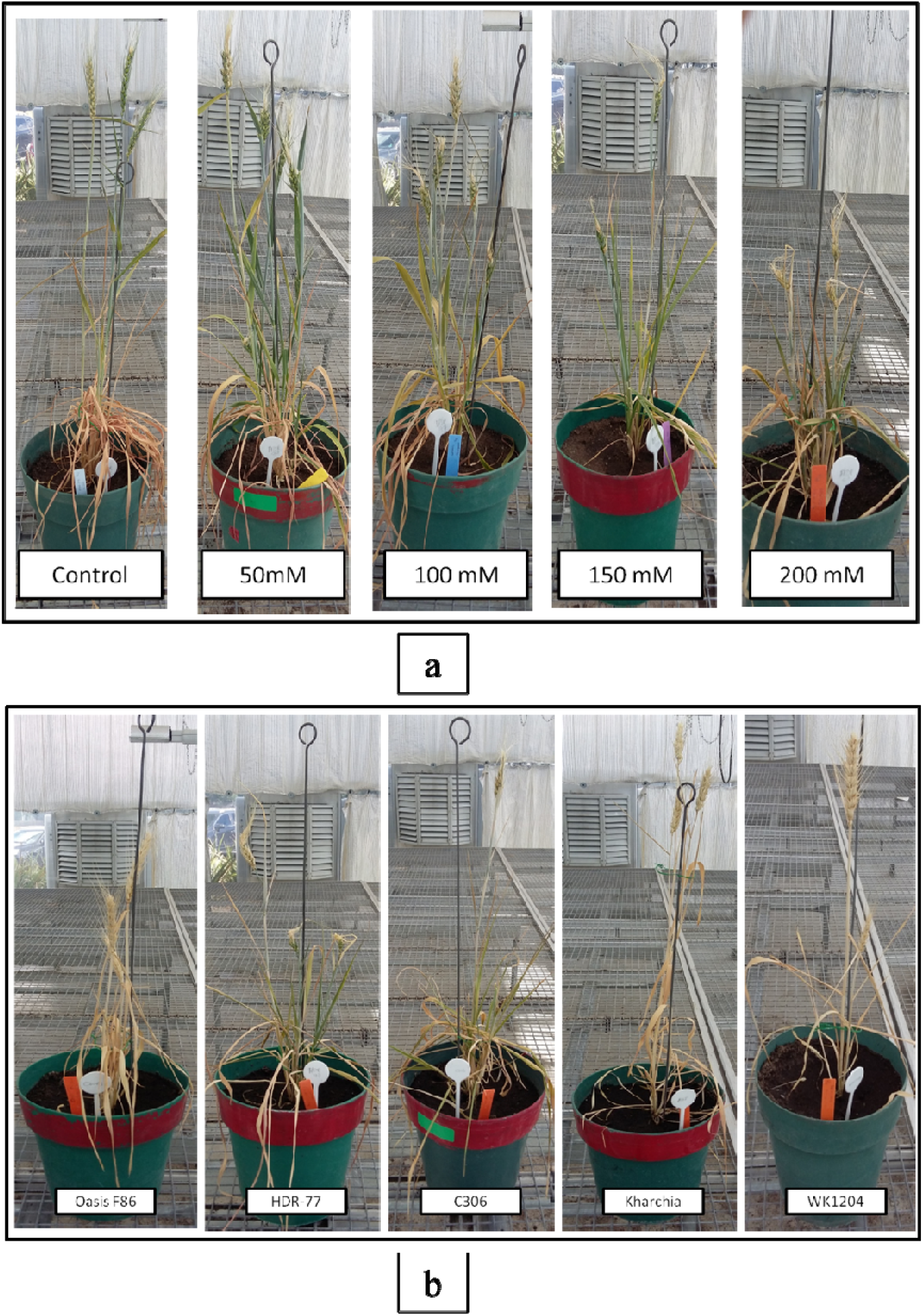
a) Salinity induced growth retardation of the cultivar HDR-77. b) Phenotypic effect of high salinity stress with 200mM on bread wheat cultivars.

Our greenhouse experiment demonstrates that the varieties C306 and HDR-77 even shown slightly better salt tolerance over known salinity tolerant reference line Kharchia. However, wheat variety like Munal#1 evinced slightly better tolerance with germination related salinity stress than varieties WK1204, OasisF86 and Dharwar Dry. Germination related study shewed that the variety WK1204 is slightly better salinity tolerant than the variety OasisF86, while the green house experiment is contrary, as OasisF86 exhibits slightly better phenotypic features with salinity stress than the variety WK1204. Perhaps, a known saline sensitive variety OasisF86 is very sensitive during germination with salinity stress. Nonetheless, the varieties C306 and HDR-77 had better tolerance with greenhouse study and germination related salt stress study (Table 5, Table 6, Supplementary Table 7).

## DISCUSSION

To better understand the sequence variation of *TaHKT1;5-D* gene in bread wheat, a gene fragment of about 3 Kb was sequenced in bread wheat varieties, while a small fragment of about 1 Kb was analyzed in wild species. In bread wheat, only three SNP were identified in the 3 Kb sequences, indicating a high *TaHKT1;5-D* gene sequence conservation within this species. However, a lot of of inter-specific SNPs was found when comparing the gene sequences of bread wheat and wild relatives. In congruence with previous studies (Huang *et al*., 2008), *HKT1;5* gene could not be amplified in *T. urartu*, supporting the conclusion that the gene might be absent in the A^u^ genome donor. Thus, none of the variant of *HKT1;5* gene was observed in A genome of wheat (Huang *et al*., 2008). However, *TmHKT1;5-A* from *T. monococcum* was presumed as strong candidate gene for *Nax2* locus, which is homeolog to *Kna1* locus with a single variant located at distal 5AL chromosome in bread wheat (Byrt *et al*., 2007). Being an allogamous species and B genome progenitor, *A. speltoides* tends to have slightly higher mutations/variant form of gene, thus we could achieve amplification only in some accessions. Hence specific attention must be given in this species to excavate the hidden form of this gene to answer whether it is due to supernumerary nature of B chromosomes with more intraspecific nucleotide variations (Jones et al., 2008). Also, this kind of study would provide better understanding of orthologous gene variations and gene flow of wild relatives and domesticated species. Gene flow from wild to domesticated species through pre-breeding entails the conservation of genetic diversity (Papa *et al*., 2005) and alien transfer of genes with functional relevancy (Jauhar *et al*., 2003).

Phylogenetic analysis among the orthologs of *HKT1;5* from diverse species of Poaceae revealed the existence of five groups based on the closeness of each species or chromosomal relevancies. The existence of multiple variants in putative B genome donor *A. speltoides*, would provide a clue for the presence of 3 variants in 4BL chromosome (Huang *et al*, 2008) of bread wheat. However, the *A. speltoides* allele 1 and 2 (from ortholog group II), revealed the similarity with the gene from 4DL chromosome and exhibited the vicinity to *T. monococcum* highlights the homeolog comparisons of *Nax2* locus and *Kna1* locus. Further comparison of ortholog group III and IV, revealed the presence of maximum variants in B genome itself or B genome related sitopsis section species *A. sharonensis*. Further the ditelosomic line experiments revealed that one homeolog is exclusively present on 4DL chromosome and the other one on 4BL of bread wheat, the possibility that this homeolog region from 4BL could have ancestrally translocated to 4DL through homoeologous recombination after AABB+DD polyploidization event, perhaps any mutation at chromosome 5B encoded mutant *Ph1* locus would have not suppressed the homoeologous recombination (Koo et al., 2020, Gaeta et al., 2009, Qi et al., 2007) or *Ph1-inhibiting gene* called *Ph*^*1*^ kind of gene would have rendered homoeologous pairing/recombination by inhibiting *Ph1* gene (Jauhar 2003). A recent study with resequencing of diploid progenitors of AADD allotetraploid line revealed the similar trend about the nonreciprocal homeologous exchange between 4A and 5D (Zhang et al., 2020) on the regions of ancient translocations of long arms between chromosomes 4A and 5A in *T. urartu* (Ling et al., 2018).

Several previous genetic analyses also compared the monocots and dicots using different genes and genomic data (Subburaj *et al*., 2016, Sarma *et al*., 2014, Cserhati., 2015). It is noteworthy that the model plant *A. thaliana* and its close relatives from family *Brassicaeae* reportedly represented a vital role in evolution and functional relevancies of plant kingdom (Koornneef and Meinke *et al*., 2010; Hu *et al*., 2011).

A comprehensive review on physiological and genomics aspects concerning the *HKT* gene between *Arabidopsis* and monocots was also reported (Horie *et al*., 2009). Besides, our study concerning the alignment of the proteins *AtHKT1* and *TaHKT1;5-D*, disclosed the identity of numerous amino acids and conservative substitution of amino acids. Secondary structure and physio-chemical properties also revealed the close vicinity between the two proteins. Homology modelling based tertiary structure prediction exhibited a least structural change between these orthologous proteins. The comparison of the presence of negatively charged and positively charged amino acids were closer. The instability index was below 40, stating that the stability of both proteins was not affected. Therefore, there was no maximum structural change or dissimilarity observed between these orthologous proteins.

The remnants of SINE9-OS and TABARE (Retroviral like sequence of plants) were observed in 5’ UTR region of the *TaHKT1;5* gene. The exon1 was disrupted with remnants of LTR retrotransposon Copia-1_SSa-I, EnSpm-14_SBi and RTE-11_BF autonomous Non-LTR Retrotransposon. Exon 1 and exon 2 was separated by a complete non-autonomous mariner DNA transposon DNA-2-N2_TA, which occupied almost all intron1. Intron 2 neither produced any blast hits on repeated elements nor showed any transposon signals/signatures. The alignment of the mRNA region (exon 2 and exon 3) from *TaHKT1;5* gene with *A. thaliana HKT1* mRNA, resulted a clear donor and acceptor sites alignment between these two orthologues genes. *A. thaliana HKT1* gene (derived from chromosome 4: Genbank CP002687, Mayer *et al*., 1999) showed higher number of transposon remnants including DNA and Retrotransposons than the gene of *TaHKT1;5*. Therefore, the size of the gene was modulated with 3680 bp and 2087 bp (from start to stop codon) in *A. thaliana* and *T. aestivum*, respectively. Unlike wheat, only one *HKT1* gene was reported in *A. thaliana* (Uozumi *et al*., 2000) and may have been diverged from its ancestor and evolved substantially in *in* Poaceae members of monocots due to the presence of several paralogs (Waters et al., 2013). Global alignment between the gene of *AtHKT1* and *TaHKT1;5-D*, revealed higher conservation/similarity in CDS region from both species. However, the introns showed least conservation, for instance remnant of mariner transposon in intron1 of *AtHKT1* showed similarity in *TaHKT1;5-D* with same region. BLASTN analysis showed that the existence of this mariner element in almost all chromosomes of bread wheat at 3120 BLASTN hits with >75% sequence similarity from A, B, D genomes of about 1000 hits per genome. Whereas 740 BLASTN hits in *A. tauchii*, 1068 and 1030 hits in *T. urartu* and *A. speltoides* respectively with >75 % sequence similarity was observed. It appeared that after polyploidization, the D genome reinforced with the multiple movements of this transposon as D genome progenitor species *A. tauschii* exhibited lesser hits for this element than D genome in bread wheat. The BLASTN analysis from the survey sequences (IWGSC-URGI and NCBI wheat) revealed that the entire gene was present in Chromosome 4DL of bread wheat. Chromosome 4BL also found to contain entire length of the gene as reported by Huang *et al*., 2008. *A. speltoides* resulted the presence of multiple variants of the gene in its genome. Any of the A chromosomes did not show reasonable query coverage, although remnants of the gene (region belongs to non-autonomous mariner transposon corresponding to intron 1) present in almost all chromosomes even from chromosome A. It is important to note that BLASTN analysis revealed the presence of remnants of various parts of the gene (for instance the region of exon 2 and intron 2 of about 20 %) in *T. urartu*, but not the whole gene. However, the presence of the gene was already reported in *T. monococcum* and *A. tauschii* (Byrt *et al*., 2007) and BLASTN hits confirmed the same. The presence of the gene in *T. durum* but absence in *T. urartu*, suggested that during or after polyploidization, it could have inherited from B genome donor rather than A genome donor *T. urartu* (Huang *et al*., 2008).

Salinity tolerance assessment was made during the germination of drought and heat tolerant cultivars such as HDR-77 (Allele ‘C’: g.[830T>C;832T>C;2106;C>T]) and C306 (Allele ‘C’) respectively and salt sensitive cultivar Oasis F86 (Allele ‘C’) along with an unknown cultivar for salinity tolerance namely WK1204. The bread wheat variety HDR-77 was reported to have higher water-binding characteristics and seed membrane stability, which confers increased drought tolerance (Chatterjee and Nagarajan, 2006). The cultivar C306 is a traditionally known drought and heat tolerant variety, which is being extensively used for the cultivation in rain-fed areas. The varieties WK1204 and Oasis F86 are known to yellow rust resistance (Thapa D.B. 2020) and saline sensitiveness (Kazi and Hettel, 1995) respectively. Our study revealed that saline tolerant cultivars were able to elongate their radicle length more than the sensitive cultivars, while the radicle numbers were less in tolerant cultivars than the sensitive cultivars. Saline sensitive cultivars were able to produce a greater number of radicles than the tolerant ones and it is contrast to radicle length, similar study was previously reported (Rahman et al., 2016). Green house study revealed that generally abiotic stress tolerant cultivars are tolerant to salinity as well, even the drought or heat tolerant varieties such as HDR-77 or C306 showed better salinity tolerance and exhibited increased yield related parameters. The prevalent ‘C’ allele is observed in more lines especially in tolerant lines, while the allele ‘A’ was specific to WK1204. Our study revealed that varieties Dharwar Dry and WK1204 are saline sensitive and comparable to reference salinity sensitive line Oasis F86, which possess allele ‘C’. Few wheat lines observed with, heterozygous forms such as Dharwar Dry with ‘B’,’C’ and Kharchia 65 with ‘C’’D’. We presume that homozygous form of ‘A’ allele is a rare form of allele only present in WK1204, while at heterozygous form ‘B’ and ‘D’ allele were observed in Dharwar Dry and Kharchia 65 lines along with their ‘C’ allele.

The most common allele ‘C’ was present both in saline tolerant and sensitive cultivars, while Allele ‘A’, ‘B’ and ‘D’ were rare and observed with both tolerant and sensitiveness depends upon the cultivars. The current study has deciphered the most redundant allele in both tolerant and sensitive cultivars that is allele ‘C’, however additional study is needed to understand why this allele is widely present among the studied cultivars. However, some cultivars were observed other alleles or combination of alternative form of gene with heterozygotic nature. Therefore, there is a need of additional studies on various genes linked through metabolic pathways or even transcription factors. Nonetheless, the current study provided the presence of multiple variants or alleles in wheat and progenitor species and the plausible comparison with phenotypes during salinity exposure. Overall, the current study proved the salinity tolerance and sensitiveness in known tolerant, sensitive and unknown lines of bread wheat and compared with *HKT1;5* alleles. However, salinity tolerance is complex physiological and genetic trait, hence further studies such as breeding and pre-breeding strategies are necessary to develop a better tolerant variety with sustainable yield. Hence, we conclude that salinity tolerance is a complex form of physiological trait and further attention with genetic focus for the validation of other interacting genes and factors of salinity tolerance is needed. Nevertheless, our study on SNPs/allele mining, phenotypic evaluation with possible correlation of alleles in wheat with an evolutionary focus would be an asset to develop a sustainable salinity tolerant wheat variety with marker assisted selection and breeding programs.

## CONCLUSION

This study proved the presence of multiple variants of the *HKT1;5* gene in bread wheat and its wild relatives. The paralogous/orthologous variants or major alleles exhibited number of SNPs variation, while minor alleles showed lesser SNPs variations. The alleles comparison with salinity phenotypes provided the evidence of allelic variants of *HKT1;5* gene behind in possible salinity tolerance and sensitiveness, however further validation is needed. SNPs/alleles and ortholog/paralog variants identified from this study, can be extensively used at intra or inter-specific breeding to enhance the salinity tolerance in bread or durum wheat, upon introgression of this gene from the compatible wild relatives. Conservative nature of this gene between monocots and dicots was also elucidated. Allele mining and evolutionary focus would be an asset for pre-breeding strategies for conservation of genetic diversity and genetic resources with an alien transfer of genes to excavate enigmatic properties of wild relatives. Overall, these analyses highlight the presence of multiple allelic variants in wheat as well from their progenitors from different geographical origins.

## Supporting information

HKT_supplementary file_Rev3

## ACKNOWLEDGEMENT

Authors duly acknowledge the research support by Bioversity International’s Vavilov-Frankel Fellowship, Italy, funded by the Grain Research Development Corporation (GRDC), Australia. Authors are grateful to Dr. Masahiro Kishii (CIMMYT) for providing ditelosomic wheat lines for this study. Authors wish to thank Dr. Susanne Dreisigacker of Biosciences Department at CIMMYT, Mexico for laboratory access to carryout molecular biological experiments. Authors also wishes to express their sincere gratitude for the encouragement from Prof. Mario Enrico Pè, Scuola Superiore Sant’ Anna, Italy, Dr. Bal Krishna Joshi, Dr. Bharat Adhikary and colleagues of CIMMYT, Mexico, ENEA, Italy and NARC, Nepal for their constant encouragement and support.

## AUTHOR CONTRIBUTIONS

Devised and designed the experiments: KT, PM, AKJ. Performed the experiments, data generated, analysed, and wrote the manuscript: KT. Assisted in manuscript revisions: PV, DS, RV, AKJ, PM. Critical suggestions for manuscript improvement: PV, HR, GV, PG, AL, CC, EP. Contributed reagents/materials/analysis tools: KT, PM, AKJ, DT, SP.

## DECLARATION

The authors have declared no competing interests.

