## Supplementary material for "Allele mining, evolutionary genetic analysis of *TaHKT1;5* gene and evaluation of salinity stress in selected lines of wheat": HKT_supplementary file_Rev3

**Supplementary Table 1.** List of primers used to amplify the alleles, orthologous and paralogous forms of *HKT1;5* gene in wheat and its wild relatives. D genome specific and A genome specific primers shown with bold letter at end of the primer name and shadowed with the genome specific base at 3’ end. Other primers are common internal primers.

| **S.No** | **Primer Sequence 5’ to 3’** | **Name** | **Tm °C** |
| --- | --- | --- | --- |
| 1 | GCCTGAAATGAATGCAAGAAAA**CA** | 24HF**A** | 67.8 |
| 2 | GCCTGAAATGAATGCAAGAAAA**TG** | 24HF**D** | 67.2 |
| 3 | ATGAGGTACTCGGCATAATGAAAATATGTTCAG**G** | 34HR**A** | 72.3 |
| 4 | ATGAGGTACTCGGCATAATG**A** | 21HR**A** | 60.9 |
| 5 | GAGGTACTCGGCATAATGGAAATATGTTCAG**A** | 32HR**D** | 71.5 |
| 6 | GAGGTACTCGGCATAATG**G** | 19HR**D** | 59.5 |
| 7 | AACATCAACGGACAAATTTTTACA | HKT5’UTRF | 63.0 |
| 8 | TGGGGGTTGGAGAAGGAC | HKTE1R | 65.1 |
| 9 | AAGTTGAGGGGGTCATCG | HKTE2R | 62.6 |
| 10 | GCATGGATGACGTCGATC | HKTE1F | 62.9 |
| 11 | AGGCACACCGGTGAGATG | HKTE1F2 | 65.1 |

**Supplementary Table 2.** Distant matrix indicating the distant index among bread wheat alleles and major alleles of *T. monococcum and* *A. tauschii.*

|  | 1 | 2 | 3 | 4 | 5 | 6 | 7 | 8 | 9 | 10 | 11 | 12 | 13 | 14 | 15 | 16 |
| --- | --- | --- | --- | --- | --- | --- | --- | --- | --- | --- | --- | --- | --- | --- | --- | --- |
| Kharchia65_AlleleD |  | 0.00 | 0.00 | 0.00 | 0.00 | 0.00 | 0.00 | 0.00 | 0.00 | 0.00 | 0.01 | 0.01 | 0.01 | 0.01 | 0.00 | 0.00 |
| TM4_PI264935_AlleleM1 | 0.00 |  | 0.00 | 0.00 | 0.00 | 0.00 | 0.00 | 0.00 | 0.00 | 0.00 | 0.01 | 0.01 | 0.01 | 0.01 | 0.00 | 0.00 |
| WK1204_AlleleA | 0.00 | 0.00 |  | 0.00 | 0.00 | 0.00 | 0.00 | 0.00 | 0.00 | 0.00 | 0.01 | 0.01 | 0.01 | 0.01 | 0.00 | 0.00 |
| TM3_PI326317_AlleleM1 | 0.00 | 0.00 | 0.00 |  | 0.00 | 0.00 | 0.00 | 0.00 | 0.00 | 0.00 | 0.01 | 0.01 | 0.01 | 0.01 | 0.00 | 0.00 |
| Munal#1_AlleleB | 0.00 | 0.00 | 0.00 | 0.00 |  | 0.00 | 0.00 | 0.00 | 0.00 | 0.00 | 0.01 | 0.01 | 0.01 | 0.01 | 0.00 | 0.00 |
| TM5_PI272560_AlleleM1 | 0.00 | 0.00 | 0.00 | 0.00 | 0.00 |  | 0.00 | 0.00 | 0.00 | 0.00 | 0.01 | 0.01 | 0.01 | 0.01 | 0.00 | 0.00 |
| TM1_PI272558_AlleleM1 | 0.00 | 0.00 | 0.00 | 0.00 | 0.00 | 0.00 |  | 0.00 | 0.00 | 0.00 | 0.01 | 0.01 | 0.01 | 0.01 | 0.00 | 0.00 |
| TM5_PI272560_AlleleM3 | 0.00 | 0.00 | 0.00 | 0.00 | 0.00 | 0.00 | 0.00 |  | 0.00 | 0.00 | 0.01 | 0.01 | 0.01 | 0.01 | 0.00 | 0.00 |
| TM1_PI272558_AlleleM2 | 0.00 | 0.00 | 0.00 | 0.00 | 0.00 | 0.00 | 0.00 | 0.00 |  | 0.00 | 0.01 | 0.01 | 0.01 | 0.01 | 0.00 | 0.00 |
| Kharchia_AlleleC | 0.00 | 0.00 | 0.00 | 0.00 | 0.00 | 0.00 | 0.00 | 0.00 | 0.00 |  | 0.01 | 0.01 | 0.01 | 0.01 | 0.00 | 0.00 |
| At2_CWI94958SH_AlleleT1 | 0.02 | 0.02 | 0.02 | 0.02 | 0.02 | 0.02 | 0.02 | 0.02 | 0.02 | 0.02 |  | 0.00 | 0.01 | 0.01 | 0.01 | 0.01 |
| At3_CWI94956SH_AlleleT2 | 0.02 | 0.02 | 0.02 | 0.02 | 0.02 | 0.02 | 0.02 | 0.02 | 0.02 | 0.02 | 0.00 |  | 0.01 | 0.01 | 0.01 | 0.01 |
| TM4_PI264935_AlleleN1 | 0.04 | 0.04 | 0.04 | 0.04 | 0.04 | 0.04 | 0.04 | 0.04 | 0.04 | 0.04 | 0.04 | 0.04 |  | 0.00 | 0.01 | 0.01 |
| TM5_PI272560_AlleleN1 | 0.04 | 0.04 | 0.04 | 0.04 | 0.04 | 0.04 | 0.04 | 0.04 | 0.04 | 0.04 | 0.04 | 0.04 | 0.00 |  | 0.01 | 0.01 |
| TM2_PI428150_AlleleM1 | 0.01 | 0.00 | 0.00 | 0.00 | 0.00 | 0.00 | 0.00 | 0.01 | 0.00 | 0.00 | 0.03 | 0.02 | 0.04 | 0.04 |  | 0.00 |
| At_HKT | 0.00 | 0.00 | 0.00 | 0.00 | 0.00 | 0.00 | 0.00 | 0.00 | 0.00 | 0.00 | 0.02 | 0.02 | 0.04 | 0.04 | 0.01 |  |

**Supplementary Table 3**. Structural and Physico-chemical properties of ***AtHKT1*** and ***TaHKT1;5-D*** gene in *A. thaliana* and *T. aestivum.*

| **Structural and Physico-chemical properties** | **AtHKT1** | **TaHKT1;5** |
| --- | --- | --- |
| Number of amino acids | 506 | 516 |
| Alpha helix | 237 | 257 |
| Extended strand | 81 | 63 |
| Random coil | 188 | 196 |
| Theoretical Isoelectric focusing point | 8.86 | 8.9 |
| Negatively charged residues (Aspartic acid and Glutamic acid) | 40 | 38 |
| Positively charged residues (Arginine and Lysine) | 48 | 44 |
| Instability Index | 34.04 | 29.75 |
| Aliphatic index | 106.42 | 103.62 |

**Supplementary Table 4.** Mean values and variability of traits under different treatments of salinity stress including control and mean values and variability of each genotype with underlying phenotypic features responding to treatment. Letters not sharing similarity are statistically significant to LSD in each row at P=0.05 level. FGP= Final Germination Percentage, GE= Germination Energy, SVI= Seed Vigor Index, CL= Coleoptile Length, RL= Radicle Length, NoR= Number of radicles, ColL- Leaf Length from Coleoptile, FW= Fresh Weight, DW= Dry Weight, RWC= Relative Water Content. Statistical significance: .p<1, *P<0.05, **p<0.01, ***P<0.001, Ns- Not significant.

|  | **Treatment** | | | | | |  |
| --- | --- | --- | --- | --- | --- | --- | --- |
| **Traits** | **Control** | **50 mM** | **100 mM** | **150 mM** | **200 mM** | **LSD at P=0.05** | **F test significance** |
| **FGP** | 93.77a | 92.88a | 86.66a | 86.22a | 73.33b | 6.97 | 2.52e-06 *** |
| **GEP** | 92.44a | 91.55a | 86.66a | 85.77a | 71.11b | 9.6 | 5.25e-08 *** |
| **SVI** | 6456.55a | 5054.97b | 3195.64c | 1985.77d | 859.55e | 913.84 | < 2e-16 *** |
| **CL** | 482.33a | 478.80a | 301.26b | 152.93c | 45.86c | 112.6 | < 2e-16 *** |
| **RL** | 683.09a | 539.20b | 358.93c | 229.66d | 109.73e | 73.7 | < 2e-16 *** |
| **NoR** | 75.06a | 63.60b | 51.80c | 45.93c | 29.133d | 9.49 | < 2e-16 *** |
| **ColL** | 958.06a | 529.73b | 161.80c | 42.53c | 0.73c | 213.41 | <2e-16 *** |
| **FW** | 1245.26a | 1187.00ab | 1034.60ab | 902.40b | 890.06b | 306.5 | 0.00217 ** |
| **DW** | 536.46a | 528.26ab | 527.86ab | 494.73b | 434.80c | 39.27 | 1.68e-10 *** |
| **RWC** | 62.36a | 55.95ab | 46.00bc | 38.49c | 37.95c | 15.32 | 1.71e-05 *** |
|  |  |  | **Genotypes** |  |  |  |  |
| **Traits** | **Kharchia** | **Kharchia65** | **OasisF86** | **DharwarDry** | **DWK26** | **LSD at**  **P=0.05** | **F test Significance** |
| **FGP** | 94.22a | 93.77a | 82.66b | 82.22b | 80.00b | 10.65 | 9.78e-05 *** |
| **GEP** | 94.22a | 92.88a | 81.33b | 80.88b | 78.22b | 9.6 | 1.93e-06 *** |
| **SVI** | 4920.93a | 3555.06 b | 3175.79b | 3160.31b | 2740.40b | 913.84 | 2.22e-08 *** |
| **CL** | 442.73a | 401.40a | 223.86b | 214.80b | 178.40 b | 112.6 | 1.7e-10 *** |
| **RL** | 513.33a | 366.13b | 361.80b | 356.09b | 323.26b | 73.7 | 1.35e-09 *** |
| **NoR** | 63.93a | 61.00a | 49.33b | 47.80b | 43.46b | 9.49 | 1.34e-08 *** |
| **ColL** | 433.46a | 365.00a | 308.13a | 293.73a | 292.53a | 213.41 | 0.254 Ns |
| **FW** | 1299.60a | 1266.80a | 1055.93ab | 851.93b | 785.06b | 306.5 | 2.87e-06 *** |
| **DW** | 636.46a | 595.13b | 581.06b | 368.86c | 340.60c | 39.27 | < 2e-16 *** |
| **RWC** | 53.37a | 51.52a | 46.97a | 46.46a | 42.44a | 15.32 | 0.258 Ns |

**Supplementary Table 5.** Mean values and variability of traits under different treatments of salinity stress including control and mean values and variability of each genotype with underlying phenotypic features responding to treatment. Letters not sharing similarity are statistically significant to LSD in each row at P=0.05 level. FGP= Final Germination Percentage, GE= Germination Energy, SVI= Seed Vigor Index, CL= Coleoptile Length, RL= Radicle Length, NoR= Number of radicles, ColL- Leaf Length from Coleoptile, FW= Fresh Weight, DW= Dry Weight, RWC= Relative Water Content, MT= Mean Germination Time, CVt= Coefficient Variation of Germination Time, MR=Mean Germination Rate, U= Uncertainty of the Germination Process, Z=Synchrony. Statistical significance: .p<1, *P<0.05, **p<0.01, ***P<0.001, Ns- Not significant.

| **Traits** | **Treatment** | | | | | | **F test significance** |
| --- | --- | --- | --- | --- | --- | --- | --- |
|  | **10ml H_2_O** | **10ml 200mM** | **15ml 200mM** | **20ml 200mM** | **25ml 200mM** | **LSD at P=0.05** |  |
| **FGP** | 81.11a | 78.88a | 77.77a | 73.33a | 72.22a | 17.17 | Ns |
| **GE** | 7.1a | 5.0b | 6.0b | 3.5b | 3.5b | 1.75 | 0.000545 *** |
| **SVI** | 665.38a | 360.55b | 357.60b | 342.54b | 342.21b | 181.63 | 7.59e-06 *** |
| **CL** | 25.82a | 25.31a | 24.19a | 24.16a | 20.05a | 7.64 | Ns |
| **RL** | 77.13a | 49.31b | 45.01b | 44.75b | 43.11b | 14.37 | 3.99e-08 *** |
| **NoR** | 5.29a | 3.94b | 3.74b | 3.64b | 3.37b | 1.158 | 0.000161 *** |
| **ColL** | 57.25a | 37.08ab | 35.27ab | 33.60b | 29.1b | 23.19 | 0.0092 ** |
| **FW** | 532.77a | 524.77a | 501.222a | 497.44a | 480.44a | 60.79 | 0.0898 . |
| **DW** | 424.44a | 421.66a | 421.22a | 411.88a | 398.77a | 44.56 | Ns |
| **RWC** | 19.95a | 19.61a | 17.15a | 16.39a | 15.82a | 8.59 | Ns |
| **MT** | 2.38a | 2.03a | 1.9a | 1.7a | 1.5a | 1.06 | Ns |
| **CVt** | 17.15a | 14.72a | 13.78a | 8.71a | 8.02a | 9.71 | 0.032301 * |
| **MR** | 0.75a | 0.72a | 0.59a | 0.54a | 0.51a | 0.34 | Ns |
| **U** | 1.87a | 1.54ab | 1.23ab | 1.14ab | 0.49b | 1.32 | 0.0481 * |
| **Z** | 2.44a | 0.77a | 0.66a | 0.25a | 0.22a | 3.07 | Ns |
|  |  |  | **Genotypes** |  |  |  |  |
|  | **Traits** | **Munal#1** | **WK1204** | **DWK26** | **LSD at P=0.05** | | **F test significance** |
|  | **FGP** | 96.66a | 86.00a | 47.33b | 11.16 |  | 5.81e-13 *** |
|  | **GE** | 5.8a | 5.4a | 2.8b | 1.35 |  | 1.02e-05 *** |
|  | **SVI** | 577.40a | 451.33b | 212.24c | 118.13 |  | 1.44e-08 *** |
|  | **CL** | 27.4a | 22.7ab | 21.63b | 4.97 |  | 0.0142 * |
|  | **RL** | 59.03a | 53.16a | 43.39b | 9.34 |  | 0.000673 *** |
|  | **NoR** | 4.24a | 4.01a | 3.74a | 0.75 |  | Ns |
|  | **ColL** | 38.93a | 38.30a | 38.15a | 15.08 |  | Ns |
|  | **FW** | 558.26a | 492.06b | 471.66b | 39.54 |  | 8.08e-06 *** |
|  | **DW** | 458.53a | 399.73b | 388.53b | 28.98 |  | 8.15e-07 *** |
|  | **RWC** | 18.05a | 17.74a | 17.56a | 5.58 |  | Ns |
|  | **MT** | 2.41a | 1.78ab | 1.63b | 0.69 |  | 0.0175 * |
|  | **CVt** | 18.19a | 11.82b | 7.41b | 6.31 |  | 0.000561 *** |
|  | **MR** | 0.73a | 0.66ab | 0.47b | 0.22 |  | 0.0184 * |
|  | **U** | 1.54a | 1.28a | 0.95a | 0.86 |  | Ns |
|  | **Z** | 1.59a | 0.82a | 0.20a | 1.99 |  | Ns |

**Supplementary Table 6.** Observation of Booting/heading in control and salinity stress induced cultivars 50 - Days after sowing (DAS). Repli- Replication, Conc- Concentration, B- Booting stage, H- Heading Stage. Y-Yes, N-No.

|  | **Genotypes** | | | | | | | | | | | | |
| --- | --- | --- | --- | --- | --- | --- | --- | --- | --- | --- | --- | --- | --- |
|  | **Repli.** | **C306** | | **HDR-77** | | **Kharchia** | | **WK1204** | | **OasisF86** | | **DWK26** | |
| **Conc.** | **Stage** | **B** | **H** | **B** | **H** | **B** | **H** | **B** | **H** | **B** | **H** | **B** | **H** |
| **0** | 1 | N | N | N | N | N | N | Y | Y | Y | Y | Y | Y |
| **0** | 2 | N | N | N | N | N | N | Y | N | Y | Y | Y | N |
| **0** | 3 | N | N | N | N | N | N | N | Y | Y | Y | Y | N |
| **50mM** | 1 | Y | N | N | N | N | N | Y | N | Y | Y | Y | Y |
| **50mM** | 2 | N | N | N | Y | N | N | Y | N | Y | Y | Y | N |
| **50mM** | 3 | Y | N | N | N | N | N | Y | N | Y | Y | Y | N |
| **100mM** | 1 | N | N | N | N | Y | N | Y | N | Y | Y | Y | N |
| **100mM** | 2 | N | N | N | N | N | N | Y | N | Y | Y | Y | N |
| **100mM** | 3 | N | N | N | N | Y | N | Y | N | Y | Y | Y | Y |
| **150mM** | 1 | Y | N | N | N | N | N | Y | N | Y | Y | Y | N |
| **150mM** | 2 | Y | N | N | N | N | N | N | N | Y | Y | Y | Y |
| **150mM** | 3 | N | N | N | N | N | N | Y | N | Y | Y | Y | Y |
| **200mM** | 1 | N | N | N | N | N | N | Y | Y | Y | Y | Y | Y |
| **200mM** | 2 | N | N | N | N | Y | N | Y | N | N | N | Y | Y |
| **200mM** | 3 | N | N | N | N | N | N | Y | N | Y | Y | Y | Y |

**Supplementary Table 7.** Alleles conferring salinity tolerance and sensitiveness from diverse wheat varieties.

| ***T. aestivum* cultivars** | **Allele** | **GenBank** | **Size** | **Salinity tolerance level** | **Current study proved** |
| --- | --- | --- | --- | --- | --- |
| *T. aestivum* WK1204 | A | KU184266 | 3073 bp | Unknown | Sensitive |
| *T. aestivum* Dharwar Dry | B | KU212871 | 2826 bp | Unknown | Sensitive |
| *T. aestivum* Munal#1 | B | KU184267 | 2438 bp | Unknown | Moderate |
| *T. aestivum* Dharwar Dry | C | KU212872 | 2826 bp | Unknown | Sensitive |
| *T. aestivum* OasisF86 | C | KU212870 | 3068 bp | Sensitive | Sensitive |
| *T. aestivum* HDR-77 | C | KU212869 | 3122 bp | Unknown | Tolerant |
| *T. aestivum* C306 | C | KU242566 | 1066 bp | Unknown | Tolerant |
| *T. aestivum* Kharchia | C | KU212875 | 3074 bp | Tolerant | Tolerant |
| *T. aestivum* Sakha8 | C | KU242567 | 965 bp | Tolerant | Unknown |
| *T. aestivum* YMI#6 | C | KU242568 | 1047 bp | Unknown | Unknown |
| *T. aestivum* Shorowaki | C | KU242569 | 965 bp | Tolerant | Unknown |
| *T. aestivum* Lu-26 | C | KU242570 | 1035 bp | Tolerant | Unknown |
| *T. aestivum* Kharchia 65 | C | KU212874 | 2883 bp | Tolerant | Tolerant |
| *T. aestivum* Kharchia 65 | D | KU212874 | 2883 bp | Tolerant | Tolerant |

**Supplementary Figure**

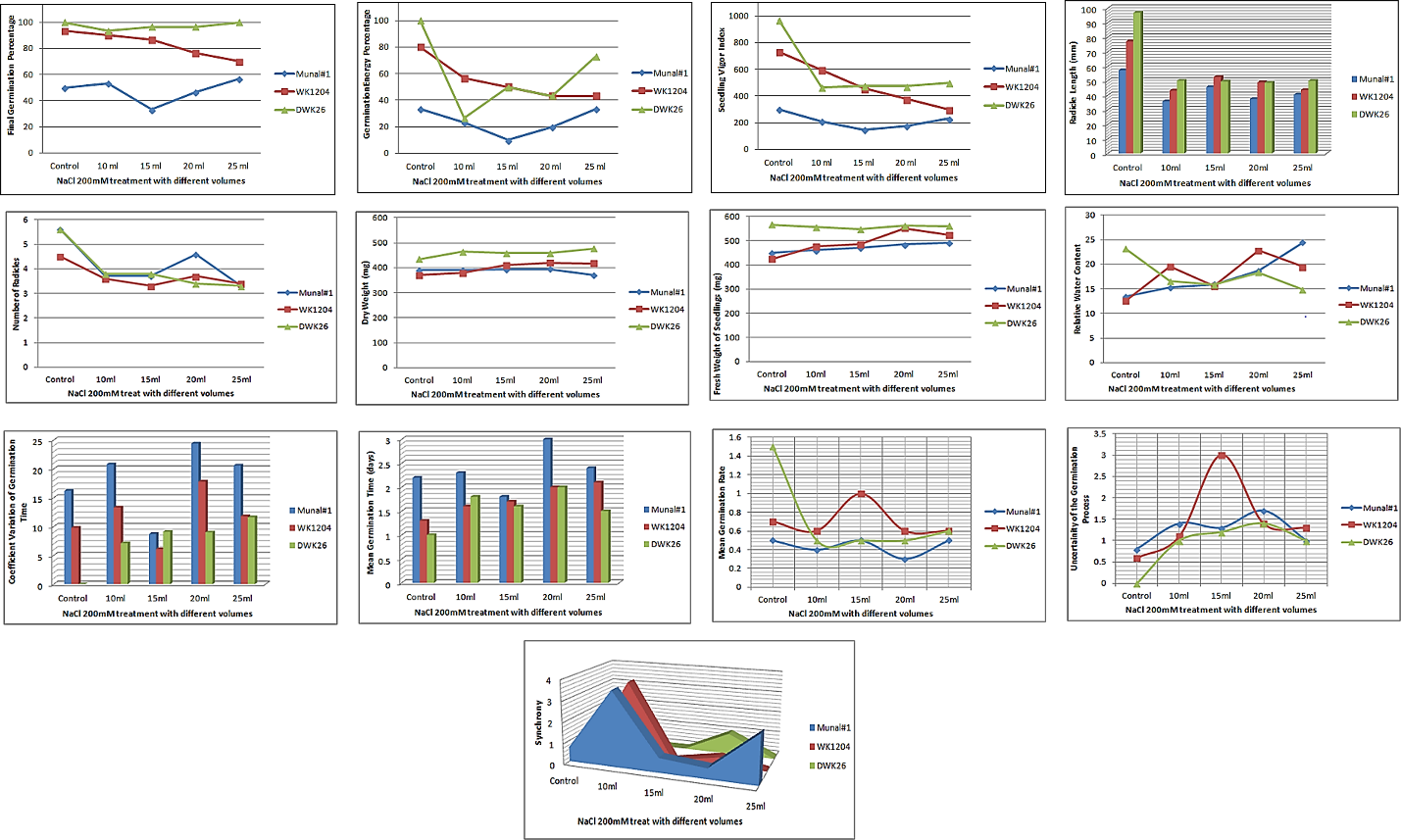

**Supplementary Figure 1.** Measured traits plotted against control and high salinity concentration (200mM) of different volumes (10ml, 15ml, 20ml, 25ml). Cultivars used: Bread wheat Munal#1, Bread wheat WK1204, Durum wheat DWK26.
